# The lupus autoantigen La is an *Xist*-binding protein involved in *Xist* folding and cloud formation

**DOI:** 10.1101/2020.03.07.981860

**Authors:** Norbert Ha, Nan Ding, Ru Hong, Rubing Liu, Xavier Roca, Yingyuan Luo, Xiaowei Duan, Xiao Wang, Peiling Ni, Li-Feng Zhang, Lingyi Chen

## Abstract

Using the programmable RNA-sequence binding domain of the Pumilio protein, we FLAG-tagged *Xist* (<u>i</u>nactivated <u>X</u> chromosome <u>s</u>pecific transcript) in live cells. Affinity pulldown coupled to mass spectrometry was employed to identify a list of 138 candidate *Xist*-binding proteins, from which, the lupus autoantigen La (encoding gene *Ssb*) was validated as a protein functionally critical for X chromosome inactivation (XCI). Extensive XCI defects were detected in *Ssb* knockdown cells, including chromatin compaction, death of female ES cells during *in vitro* differentiation and chromosome-wide monoallelic gene expression pattern. Live-cell imaging of *Xist* RNA reveals the defining XCI defect: *Xist* cloud formation. La is a ubiquitous and versatile RNA-binding protein with RNA chaperone and RNA helicase activities. Functional dissection of La shows that the RNA chaperone domain and/or the ATP binding motif play critical roles in XCI. In mutant cells, *Xist* transcripts are unstable and misfolded. These results show that La is critically involved in XCI, possibly as a protein regulating the in-cell structure of *Xist*.

## INTRODUCTION

*Xist* RNA is a prototype long non-coding RNA (lncRNA) involved in X chromosome inactivation (XCI), a mammalian dosage compensation mechanism, in which one female X chromosome is transcriptionally silenced to balance the X-linked gene dosage between males and females (1). Upon the onset of XCI during early embryonic development, the coating of *Xist* RNA transcripts on the chromosome territory of the chosen inactive X chromosome (Xi) recruits epigenetic factors for heterochromatinization and establishes the chromosome-wide gene silencing (1). Intensive efforts have been spent in isolating *Xist*-binding proteins. Three attempts of comprehensive isolation of *Xist*-binding proteins were reported in 2015 (2-4). Chu et al. identified 81 candidate *Xist*-binding proteins (4). Using a more selective approach, McHugh et al identified a list of 10 candidate proteins (3). Meanwhile, through a more sensitive approach, Minajigi et al. generated a list of more than 700 proteins (2). 4 proteins were commonly identified by all three studies. These results generate an initial portrait of a fascinating and yet still poorly understood epigenetic machinery recruited/assembled by the *Xist* RNA. These results also illustrate the difficulty of balancing the assay’s specificity and sensitivity in profiling a complex lncRNA-protein interactome.

Advanced by breakthrough technologies in isolating lncRNA-binding proteins, the list of key proteins recruited by *Xist* is growing, which includes transcription repressors (SPEN, also known as SHARP, SMRT/HDAC1-associated repressor protein), proteins involved in N^6^-adenosine (m^6^A) RNA methylation (RBM15 and WTAP), heterogeneous nuclear ribonecleoproteins (hnRNP K and hnRNP U), histone modifiers (polycomb proteins) (2-9). It is believed that *Xist* recruits the proteins and functions as a scaffold to assemble them into a protein machinery. However, the portrait of this protein machinery remains fragmented. Many questions still remain. What is the structure of the scaffold RNA? How is the protein machinery assembled and operated? How do *Xist* transcripts spread along its host chromosome?

Here, we devised a different system for profiling the *Xist* interactome. We have previously shown that the *Xist* RNA can be efficiently tagged in live cells by the programmable RNA-sequence binding domain of the Pumilio protein (PUF: Pumilio-homology domain; PBS: PUF-binding site) (10). A transgenic cell line previously generated in the lab is a male mouse ES cell line carrying an X-linked single-copy inducible *Xist* transgene with PBSb sites fused to its 5’-ends (Fig. 1A) (10). We further engineered the cell line to have it stably express a PUFb-FLAG fusion protein (i-FLAG-*Xist*, Fig. 1A). Therefore, in this cell line, the *Xist* RNA is efficiently FLAG-tagged, which enables the usage of the highly specific and sensitive anti-FLAG antibody in the subsequent protein isolation work (Fig. 1B). Meanwhile, we engineered a negative control cell line, i-Empty, which stably expresses PUFb-FLAG but carries an empty inducible cassette (Fig. 1A). We name this method “FLAG-out”. In this study, using FLAG-out, we identified 138 candidate *Xist*-binding proteins, and further validated the lupus autoantigen La as an *Xist*-binding protein functionally critical for XCI.

**Figure 1.**
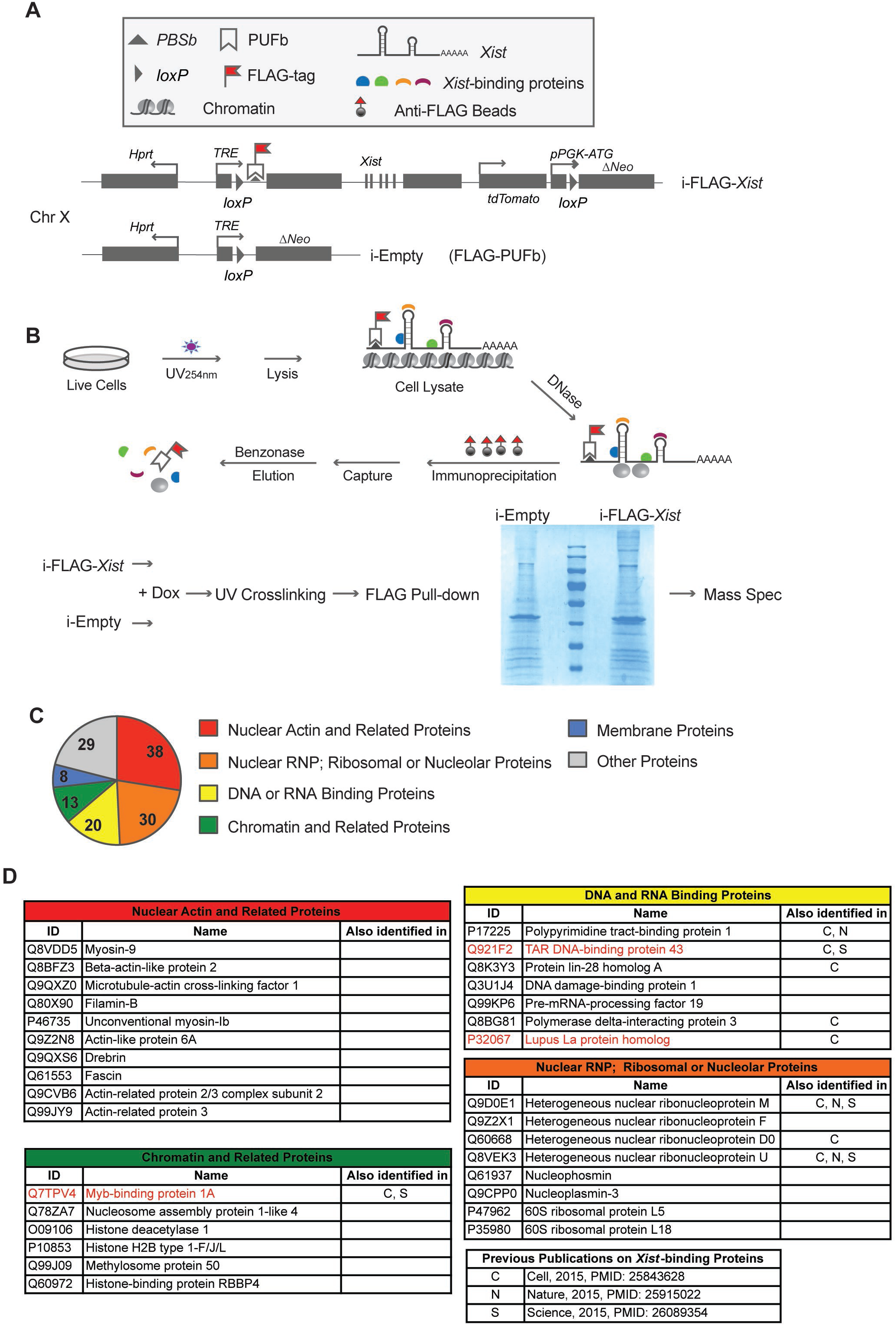
Schematic diagram of “FLAG-out” and its identified *Xist*-binding proteins. **(A)** Diagram of the transgenic mouse ES cell lines i-FLAG-*Xist* and i-Empty. **(B)** Diagram of the experimental design of “FLAG-out” and a protein gel image of the FLAG pull-down samples. **(C and D)** Candidate *Xist*-binding proteins identified by FLAG-out.

## RESULTS

### “FLAG-out” the *Xist*-binding proteins

After *Xist* induction, we fixed the cells by UV-crosslinking and performed FLAG affinity pull-down (Fig. 1B). Mass-spec was used to analyze two protein pull-down samples, a Dox-treated i-FLAG-*Xist* sample and a Dox-treated i-Empty sample (Fig. 1B). 69 proteins were identified only in i-FLAG-*Xist* but not in i-Empty (Table S2 A-C). Furthermore, for the proteins identified in both samples, we ranked them according to their protein scores in each sample and calculated the ranking gains. hnRNPM, a known *Xist*-binding protein, showed a ranking gain of 11 in i-FLAG-*Xist*. Therefore, additional 69 proteins with ranking gains higher or equal to 11 were also selected as candidate proteins (Table S2D). In total, 138 candidate proteins were selected (Fig. 1C and Table S2). Among the selected candidates, 9 proteins were also identified as *Xist*-binding proteins in previous studies (2-4). Interestingly, the 138 candidate proteins can be clearly classified into 6 functional groups: [1] nuclear actin and related proteins (38 proteins); [2] chromatin and related proteins (13 proteins); [3] DNA and RNA binding proteins (20 proteins); [4] nuclear RNP, ribosomal and nucleolar proteins (30 proteins); [5] membrane proteins (8 proteins); [6] other proteins (29 proteins) (Fig. 1 C and D, and Table S1). The complete list of the candidate proteins is shown in Table S1. Selected candidate proteins are shown in Fig. 1D.

### Validation of the functional significance of the lupus autoantigen La in the induced XCI

From the candidate proteins, we shortlisted three proteins for individual validation. Myb-binding protein 1A (Mybbp1a, Q7TPV4) and TAR DNA-binding protein 43 (Tardbp, Q921F2) were selected because they are known transcription repressors (11, 12). The Lupus autoantigen La (P32067, encoding-gene name: *Ssb*) was selected because systemic lupus erythematosus (SLE) is an autoimmune disease characterized by a strikingly high female to male ratios of 9:1 (13). Moreover, its autoimmune antigen La is a ubiquitous and versatile RNA-binding protein and a known RNA chaperone (14). All the three selected candidates have also been identified as *Xist*-binding proteins in previous studies (2, 4). Moreover, the knockout of these three genes all lead to early embryonic death. *Tardbp* knockout causes embryonic lethality at the blastocyst implantation stage (15). *Mybbp1a* and *Ssb* knockout affect blastocyst formation (16, 17). Early embryonic lethality is a mutant phenotype consistent with a critical role of the mutated gene in XCI (1).

As the gene knockout of all three candidate genes causes early embryonic lethality, we chose to knock down their expression using shRNAs to validate the candidate genes’ roles in the induced XCI. The selected cell line is a male ES cell line carrying an inducible X-linked *Xist* transgene. Inducible *Xist* expression in this cell line causes cell death due to the inactivation of the single X chromosome in male cells. Massive cell death usually occurs after 4-5 days of doxycycline (Dox) treatment, which can be used as a convenient assay to assess the functionalities of XCI (Fig. 2A). We established clonal ES cell lines stably expressing the shRNAs and confirmed the shRNA knockdown efficiencies (Fig. 2 A and B). Knocking down *Ssb* (the gene encoding La) showed a negative effect on the cell growth rate (Fig. 2 C and D), consistent with the role of La as a ubiquitous RNA-binding protein involved in house-keeping functions such as tRNA biogenesis. Nonetheless, the cell survival rates show that knocking down *Ssb* significantly rescued the cells from the toxicity of induced XCI (Fig. 2 C-E). These results confirm that La is involved in induced XCI. To rule out the off-target effect of shRNAs, we used a second shRNA construct and obtained similar results (Fig. 2 C-E). In the forgoing experiments, cells were cultured as undifferentiated ES cells during the induction. We performed the Dox-induced cell death assay in differentiating ES cells and obtained consistent results (Fig. S1). We further compared the X-linked genes’ expression levels in control cells and in *Ssb* knockdown cells during the induced XCI. The expression level of 8 selected X-linked genes were measured by quantitative RT-PCR (Fig. 2 F and G). Compared to the empty vector control, higher expression levels of X-linked genes were exclusively detected in *Ssb* knockdown cells showing the impaired XCI status in *Ssb* knockdown cells. In addition, we validated the interaction between La and *Xist* RNA by RNA immunoprecipitation (RIP) (Fig. S2). These results confirm that La is involved in the induced XCI in the male transgenic cell line.

**Figure 2.**
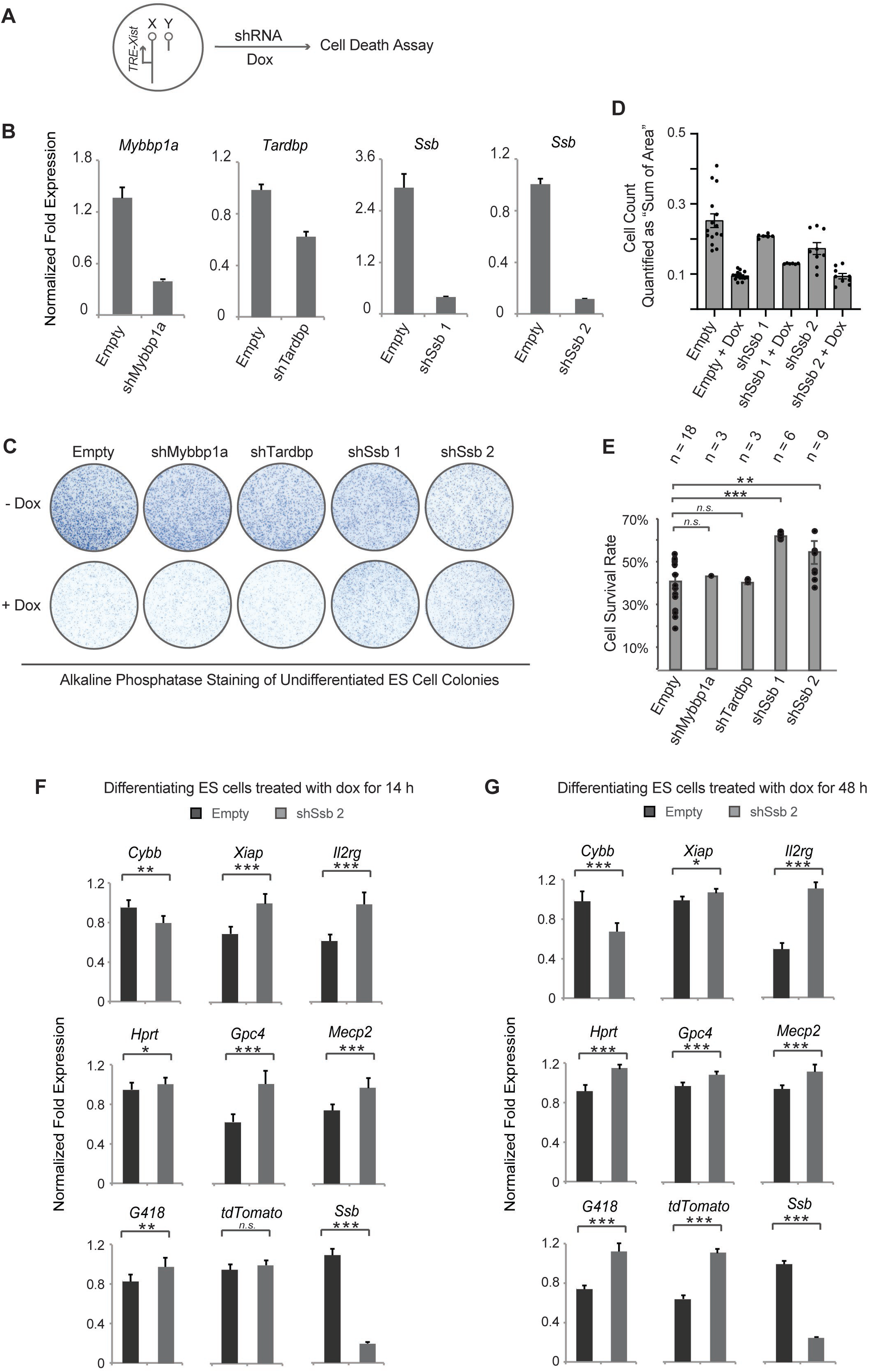
Individual validation of three candidate *Xist*-binding proteins. **(A)** The experimental design of the cell survival assay. **(B)** Quantitative RT-PCR to assess the effect of shRNA knockdown. Clonal ES cell lines stably transfected with the shRNA constructs were established. Cells stably transfected with the empty shRNA vector (Empty) serve as the control. Data are shown in relative fold expression. Normalization was performed using *Actb* and *Gapdh*. Error bars indicate SEM (n=3). **(C)** Representative macrographs of alkaline phosphatase staining on undifferentiated ES cell colonies, using EGFP-iXist cells (10). Clonal ES cell lines stably transfected with the shRNA constructs against *Mybbp1a, Tardbp, Ssb* and the empty shRNA vector were cultured as undifferentiated ES cells. Dox treatment was carried out for 5 days. **(D)** Cell count was quantified as the sum of area of alkaline phosphatase staining. Data are shown as mean ± SEM. n = 15 for Empty; n = 6 for *shSsb*1; n = 9 for *shSsb*2, from at least two independent experiments. **(E)** Cell survival rate was calculated by alkaline phosphatase-stained colony counts. Data are shown as mean ± SEM. The statistical analysis used is the Student’s *t*-test. **p < 0.01; ***p < 0.001; n = the number of experimental replications. **(F and G)** Quantitative RT-PCR to assess the effect of *Ssb* knockdown on X-linked gene expression during the induced XCI. 8 X-linked genes were selected. *tdTomato* and *G418* (neomycin selection cassette) are two X-linked transgenes and the rest are endogenous X-linked genes. *Ssb* (chromosome 2) was included to show the effect of shRNA knockdown. Of these selected genes, *Gpc4, Hprt, Mecp2, G418*, and *TomatoRed* are silenced at 24 and 48 hours after Dox induction. *Mecp2, G418*, and *TomatoRed* are silenced as early as 8 hours after Dox induction (34). Cells were cultured as differentiating ES cells and Dox treatment was carried out for 14 hours or 48 hours. Data are shown in relative fold expression. Normalization was performed using *Actb*. Error bars indicate SEM (n=6, from two independent experiments). The statistical analysis used is the Student’s *t*-test. n.s > 0.05; *p < 0.05; **p < 0.01; ***p < 0.001.

Meanwhile, our results do not confirm the functional roles of Mybbp1a and Tardbp in XCI (Fig. 2 C and E). However, it is possible that the knockdown efficiencies of these two genes were not high enough to overcome the thresholds required to disturb XCI. Alternatively, there might be some redundant factors for Mybbp1a and Tardbp that compensate the loss of these two factors.

### Validation of the functionality of La in endogenous XCI in female cells

To validate the functionality of La in endogenous XCI, we performed shRNA knockdown of *Ssb* in female ES cells (Fig. 3A). We confirmed the knockdown efficiency on the selected clonal cell lines stably expressing shRNAs against *Ssb* (Fig. S3). XCI can be triggered in female mouse ES cells by *in vitro* differentiation. Massive cell death usually occurs during this process in female mouse ES cells with XCI defects (1). Consistent with this notion, we observed significant cell death in the *Ssb* knockdown cells during *in vitro* differentiation (Fig. 3 B and C). Importantly, on the *Ssb* knockdown background, significantly more cell death was observed in female cells than male cells (Fig. 3 B and C). Although *Ssb* knockdown showed negative effect on the cell growth rate of male ES cells (Fig. 2 C and D), the morphology of male embryoid bodies (EBs) on day 4 of *in vitro* differentiation were comparable to the wild type EBs. The EBs could attach to the surface of the tissue culture flask and showed significant amount of expansion two days later (Fig. 3B). This is in clear contrast to the female EBs. With *Ssb* knockdown, the female EBs were small and unhealthy on day 4 of *in vitro* differentiation. The EBs barely attached to the surface of the tissue culture flask and showed no or very limited amount of expansion two days later (Fig. 3B). These results support a functional role of La in XCI.

**Figure 3.**
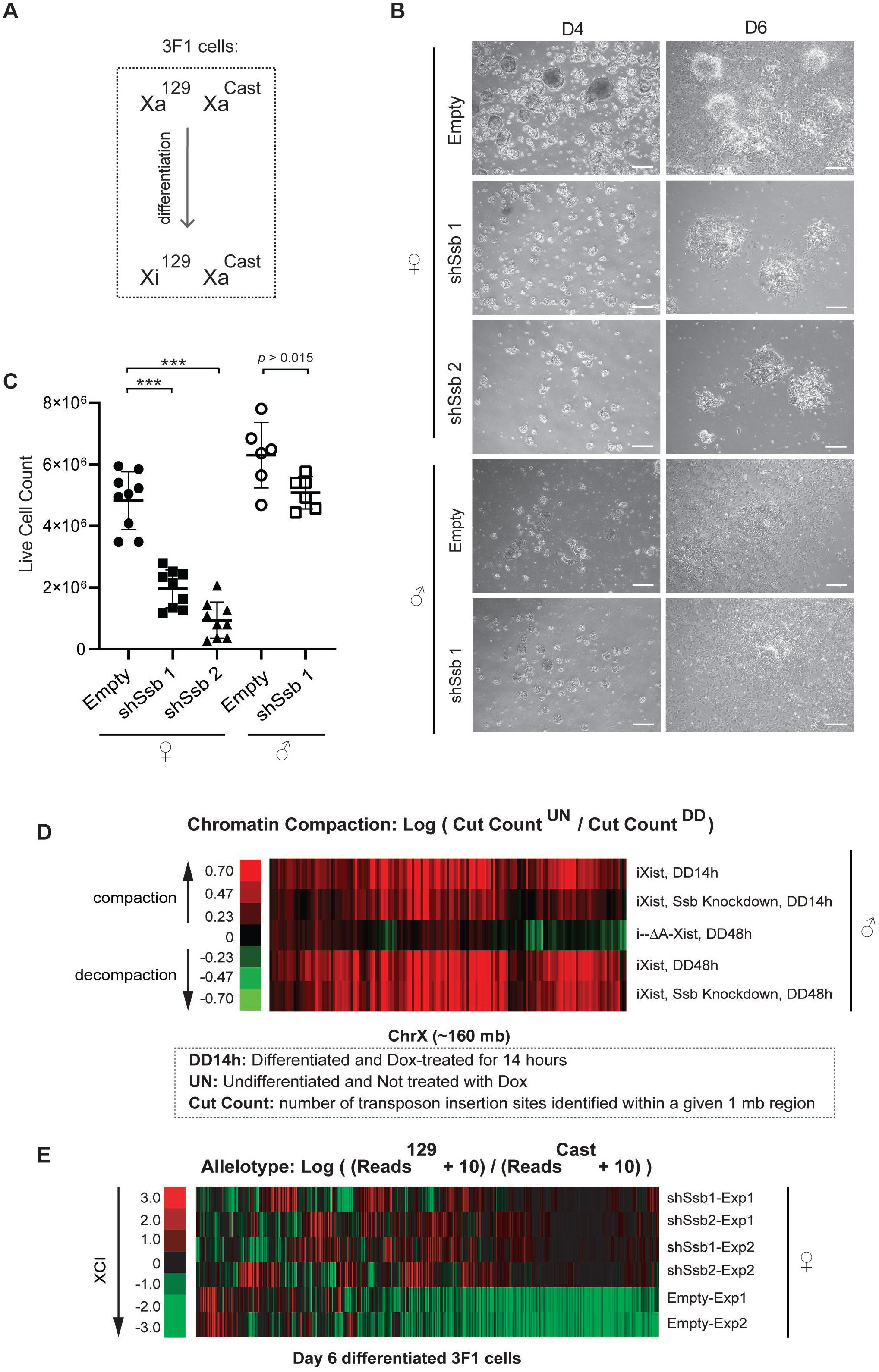
Functional validation of La in female ES cells during the endogenous XCI. **(A)** The genotype and the preemptive XCI choice phenotype of 3F1 cells. **(B)** Representative brightfield microscope images showing the differentiating 3F1 ES cells at day 4 (D4) and day 6 (D6) of *in vitro* differentiation. Scale bars, 250 μm. **(C)** Cell survival rates on *in vitro* differentiation day 6. Data are shown as mean ± SEM. The statistical analysis used is the Student’s *t*-test. ***p < 0.001; n = 9 from three independent experiments. **(D)** ATAC-seq results. The mouse X chromosome is divided into ∼160 1-mb regions. The Cut Count of each region was analyzed, and the chromatin compaction status was quantitatively measured for each region. The results are shown in heatmaps. i-ΔA-*Xist* is an inducible mutant *Xist* transgene, in which the critical A-repeat region is deleted. **(E)** Heatmaps of chromosome-wide RNA allelotyping results. RNA samples were isolated from differentiating ES cells on day 6 of *in vitro* differentiation.

As La is an RNA-binding protein involved in various house-keeping functions, the massive cell death, which occurred in *Ssb* knockdown female ES cells during *in vitro* differentiation, may not be directly and solely attributed to XCI defects. Therefore, we performed ATAC-seq (Assay for Transposase-Accessible Chromatin with high-throughput sequencing) to examine the chromatin compaction along X chromosome. We also performed padlock SNP capture to directly assess the XCI status by allelotyping the X-linked gene expression chromosome-wide.

ATAC-seq was performed using male ES cell lines carrying an X-linked inducible *Xist* transgene so that the sequencing reads mapped onto X chromosome only reflect the chromatin status of one X chromosome, the inactive X. In ATAC-seq, the density of transposon insertion reveals the status of chromatin compaction. We analyzed the transposon insertion sites genome-wide and defined “Cut Count” as the number of transposon insertion sites identified within a 1-Mb region. The chromatin compaction status during the induced XCI was then quantitatively measured by the ratio of the Cut Counts between the uninduced undifferentiated cells and the differentiated cells treated with Dox (Fig. 3D). Induced XCI clearly caused chromatin compaction in wild type cells, meanwhile the induced expression of a mutant *Xist* transgene with the critical A-repeat region deleted (i-ΔA-*Xist*) failed to generate chromatin compaction (Fig. 3D, Fig. S4, and Table S3). These results serve as controls to validate the experimental system. On the *Ssb* knockdown background, induced XCI also caused chromatin compaction, but to a lesser degree than that in wild type cells (Fig. 3D and S4). This observation was clearly made after 14 hours Dox treatment. After 48 hours Dox treatment, the chromatin compaction status was comparable between the wild type sample and the *Ssb* knockdown sample, although a slightly lesser degree of chromatin compaction was still visible in the knockdown sample (Fig. 3D and S4). These results directly connect La to the heterochromatinization of Xi, supporting a functional role of La in XCI. Meanwhile, the results also show that the *Ssb* knockdown cells encountered difficulties during the early onset of XCI. We further discuss this issue in the later sections.

To directly assess the XCI status of X-linked genes, we comprehensively allelotyped X-linked genes. The shRNA knockdown was performed on a female ES cell line 3F1 (18), which carries Xs from two genetic backgrounds, the 129 mouse strain (X^129^) and the *Mus musculus castaneus* (CAST/Ei) mouse strain (X^Cast^). The “preemptive choice” mutant phenotype of 3F1 cells causes non-random inactivation of the X^129^ allele (18) (Fig. 3A). Therefore, the XCI status of an X-linked gene can be evaluated by RNA allelotyping of X^129^ and X^Cast^, which provide ample choice of single nucleotide polymorphisms (SNPs). Padlock SNP capture, a high-throughput and high-resolution RNA allelotyping method (19), was performed to profile the XCI status of X linked genes chromosome-wide. The padlock probe library was designed to target 2,969 SNPs covering 1,110 (∼55%) of the X-linked genes (19). 457 X-linked genes were successfully allelotyped in the experiment. Padlock SNP capture detected bi-allelic expression of genes along the entire X chromosome in *Ssb* knockdown cells, demonstrating obvious XCI defects (Fig. 3E and Table S4). This result provides the direct evidence confirming the critical functionality of La in XCI.

### Knockdown of *Ssb* impairs *Xist* cloud formation

To further study the functional roles of La in the endogenous XCI, we investigated the *Xist* cloud formation in *Ssb* knockdown cells. In undifferentiated female ES cells, *Xist* expression is detected as a pinpoint signal associated with its gene locus in *cis*. Upon differentiation, *Xist* expression is up-regulated along the chosen Xi and *Xist* RNA transcripts spread out to cover the chromosome territory *in cis*. Thus, in differentiated female cells, *Xist* expression is detected as a cloud signal (the *Xist* cloud) enveloping the Xi chromosome territory. We performed *Xist* RNA FISH in differentiating ESCs. After 6 days of *in vitro* differentiation, *Xist* clouds could be detected in more than 80% of the wild type cells. However, in *Ssb* knockdown cells, *Xist* clouds were detected in a significant lower percentage of cells (Fig. 4A and S5). Given that significant cell death occurs in *Ssb* knockdown cells during *in vitro* differentiation and RNA FISH cannot be applied to dead cells, the percentage of cells showing faulty *Xist* cloud formation should be even higher in *Ssb* knockdown cells. Taken together, these results show that La is involved in *Xist* cloud formation.

**Figure 4.**
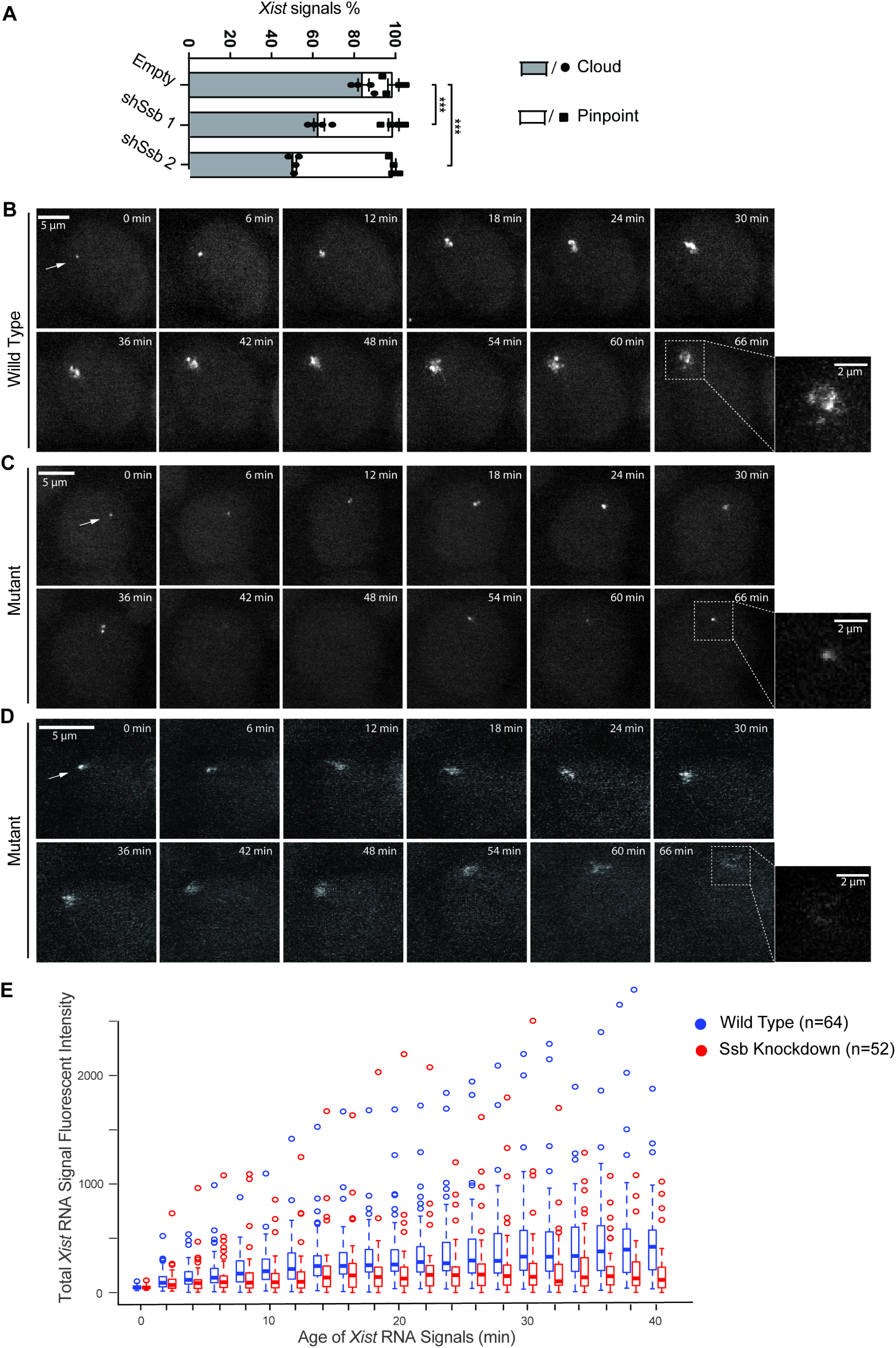
La plays critical roles in *Xist* cloud formation. **(A)** *Xist* RNA FISH results of day 6 *in vitro* differentiation female ES cells. Quantification of *Xist* RNA signals from four independent experiments is shown. Data are shown as mean ± SEM. The statistical analysis used is *X*^*2*^ test. *** p < 0.001; n = 505 for Empty; n = 540 for shSsb 1; n = 499 for shSsb 2. **(B-D)** Live-cell imaging of *Xist*. The *Xist* RNA cloud formation processes in one representative wild type cell and two *Ssb* knockdown cells are shown. Dox treatment was started after cells were *in vitro* differentiated for 16 hours. Live-cell imaging was started one hour after Dox treatment. All the live-cell imaging was carried out in a 2-hour time span with a time interval of 2 min. For direct comparison between the two *Xist* signals, images are shown in a 66 min time span with a 6 min time interval. The time point when a signal was first detected is defined as time zero for the signal. Maximum intensity z-projections are shown. **(E)** The total fluorescent intensity of an *Xist* RNA signal in live cells was measured every 2 minutes for the first 40 minutes after the signal first appeared. The measurement was done on 64 *Xist* signals from wild type cells and 52 *Xist* signals from *Ssb* knockdown cells. *Xist* signals were randomly selected from two independent experiments. The data are presented in box plots. Student’s *t-*test was performed to compare the pair of data sets at each time point. p > 0.05 for the time points of 0 – 8 min; p < 0.05 for the time points of 14 & 16 min; p < 0.009 for the rest of the time points.

To avoid the shortcomings of RNA FISH not applicable to dead cells, and to observe the process of *Xist* cloud formation during the early onset of XCI, we performed live-cell imaging of *Xist* RNA. We previously established a live-cell imaging system of the *Xist* RNA in a transgenic male mouse ES cell line carrying an X-linked single-copy inducible *Xist* transgene with PBSb sites fused to its 5’-ends (Fig. 1A). The cell line also stably expresses a PUFb-GFP fusion protein. Therefore, *Xist* RNA can be efficiently GFP-tagged in live cells. We chose an inducible transgene to study the *Xist* RNA in live cells, because the system provides a synchronized onset of XCI in the cell population. In female cells, during the *in vitro* differentiation, the onset of the endogenous XCI occurs in an unsynchronized manner. For a given cell, the onset of XCI may occur at any time during a window of a few days. However, high-quality live-cell imaging can only be performed within a 2-hour time window due to technical issues, such as photo-bleaching and phototoxicity. In our experimental system, Dox treatment was started after the cells were *in vitro* differentiated for 16 hours. Live-cell imaging was performed 1 hour after Dox treatment and lasted for 2 hours with a 2-minute time interval. Because the selected time window belongs to the early onset stage of XCI, the issue of cell death caused by XCI defects is circumvented. In wild type differentiating cells, the behavior of *Xist* cloud formation is synchronized and consistent (10). The *Xist* RNA signals first appear as small pinpoint signals that then gradually grew into ∼2 µm large *Xist* RNA clouds within 60-90 min (Fig. 4B, and Movies S1-3). In *Ssb* knockdown cells, the *Xist* cloud formation clearly encountered difficulties (Fig. 4 C-E, and Movies S4-7). In some cells, an *Xist* RNA signal first appeared as a small pinpoint signal, but the pinpoint signal failed to stabilize. Instead of growing into a large cloud signal, the pinpoint signal was on and off during the first 60-90 min after the signal first appeared (Fig. 4C, and Movies S4). In some other cells, a pinpoint *Xist* RNA signal grew into a faint cloud signal, and the cloud signal quickly diffused and vanished (Fig. 4D, and Movies S5). We analyzed 64 *Xist* signals in wild type cells and 52 *Xist* signals in *Ssb* knockdown cells. The total fluorescent intensity of each *Xist* signal was measured every 2 minutes during the first 40 minutes after the signal first appeared (Fig. 4E). The results clearly revealed the faults of *Xist* cloud formation on the *Ssb* knockdown background. Quantitative RT-PCR showed that *Ssb* depletion does not change *Xist* induction in the *Xist* inducible male ES cell line, thus excluding the possibility that *Ssb* depletion affects *Xist* cloud formation by down-regulating *Xist* RNA (Fig. S6). In summary, the observed behaviors of *Xist* cloud formation in *Ssb* knockdown cells are heterogenous but consistently show the difficulties encountered in forming the *Xist* cloud. In mutant cells, the *Xist* RNA transcripts failed to spread out and coat the X chromosome territory. It is likely that the RNA transcripts are quickly diffused and/or degraded. As *Xist* cloud formation is the initial step which triggers XCI, the defective *Xist* cloud formation should be the primary reason behind other XCI defects observed in *Ssb* knockdown cells. These results confirm that the defining XCI defect in *Ssb* knockdown cells is faulty *Xist* cloud formation.

### Knockdown of *Ssb* compromises the enrichment of Polycomb marks on Xi

Even though *Xist* cloud formation is less efficient in the mutant cells, the knockdown cell population behave heterogeneously and *Xist* clouds are formed in a substantial fraction of surviving cells. We further investigated whether La regulates other events during the later stages of XCI. It is known that, in order to inactivate one large chromosome, *Xist* cloud recruits other silencing factors to heterochromatinize Xi with multiple layers of epigenetic modifications (1). Two histone modifications enriched along Xi are the Polycomb marks, H3K27me3 and H2AK119ub (9). We performed immuno-RNA FISH to study the role of La in Polycomb marks enrichment along Xi. After 6 days *in vitro* differentiation, most of the *Xist* clouds signals in RNA FISH overlapped with enrichments of Polycomb marks detected by the immunostains (Fig. 5). However, the enrichment of Polycomb marks along Xi is significantly disrupted in *Ssb* knockdown cells. These results suggest that La is also involved in establishing the enrichment of Polycomb marks along Xi.

**Figure 5.**
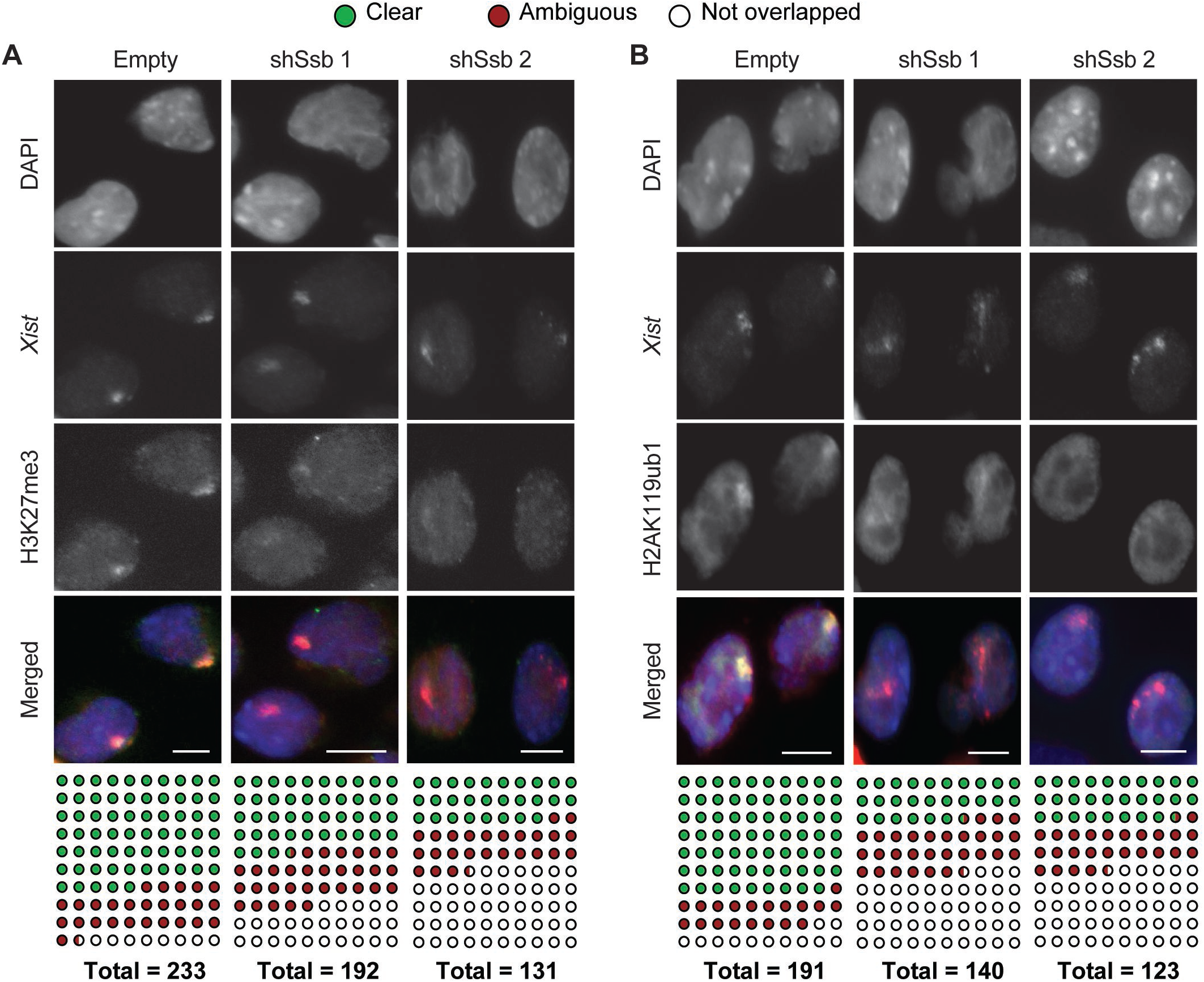
*Ssb* is involved in Polycomb mark enrichment along Xi. **(A and B)** Immuno-RNA FISH detecting H3K27me3 (A), H2AK119ub (B) and *Xist*. Female ES cells were *in vitro* differentiated for 6 days. Immunostains were performed before the RNA FISH. DNA was counter stained with DAPI (blue). Scale bars, 8 μm. The total number of *Xist* clouds with clear, ambiguous or undetectable overlapping with the histone mark enrichment were tallied and tabulated below.

### The RNA chaperone domain of La is critical in XCI

La is an RNA-binding protein with 5 defined functional domains including 3 RNA-binding domains (La motif, LAM; RNA recognition motif 1, RRM1; RNA recognition motif 2, RRM2), an RNA chaperone domain (RCD) and a nuclear localization signal (NLS) (20, 21). We generated plasmid constructs carrying in-frame deletions of the selected domains of La (Fig. 6A) and stably transfected the *Ssb* knockdown cells with these “rescue” plasmid constructs (Fig. 6B). GFP fusion was included in the plasmid design for checking transfection efficiencies and the transgene expression levels in the selected clonal cell lines (Fig. S7). We first rescued female ES cells with a full-length *Ssb* construct. Indeed, the full-length *Ssb* construct showed significant rescue effects on cell survival, *Xist* cloud formation and Polycomb marks enrichment (Fig. S8). Meanwhile, all Δ constructs showed significant rescue effects of various degrees on the cell survival during *in vitro* differentiation, except for ΔRCD and ΔNLS, indicating the critical roles of these two functional domains (Fig. 6C). For *Xist* cloud formation, ΔNLS and ΔRCD showed no rescue effect, while other Δ constructs showed significant rescue effects of various degrees (Fig. 6D). These results show that all three RNA-binding domains are required for the functionality of La in XCI and highlight the critical functions of the RCD and NLS. As XCI is a nuclear event, it is obvious why the NLS of La is critical for its functionality in XCI. On the other hand, the critical role of RCD of La in XCI is intriguing.

**Figure 6.**
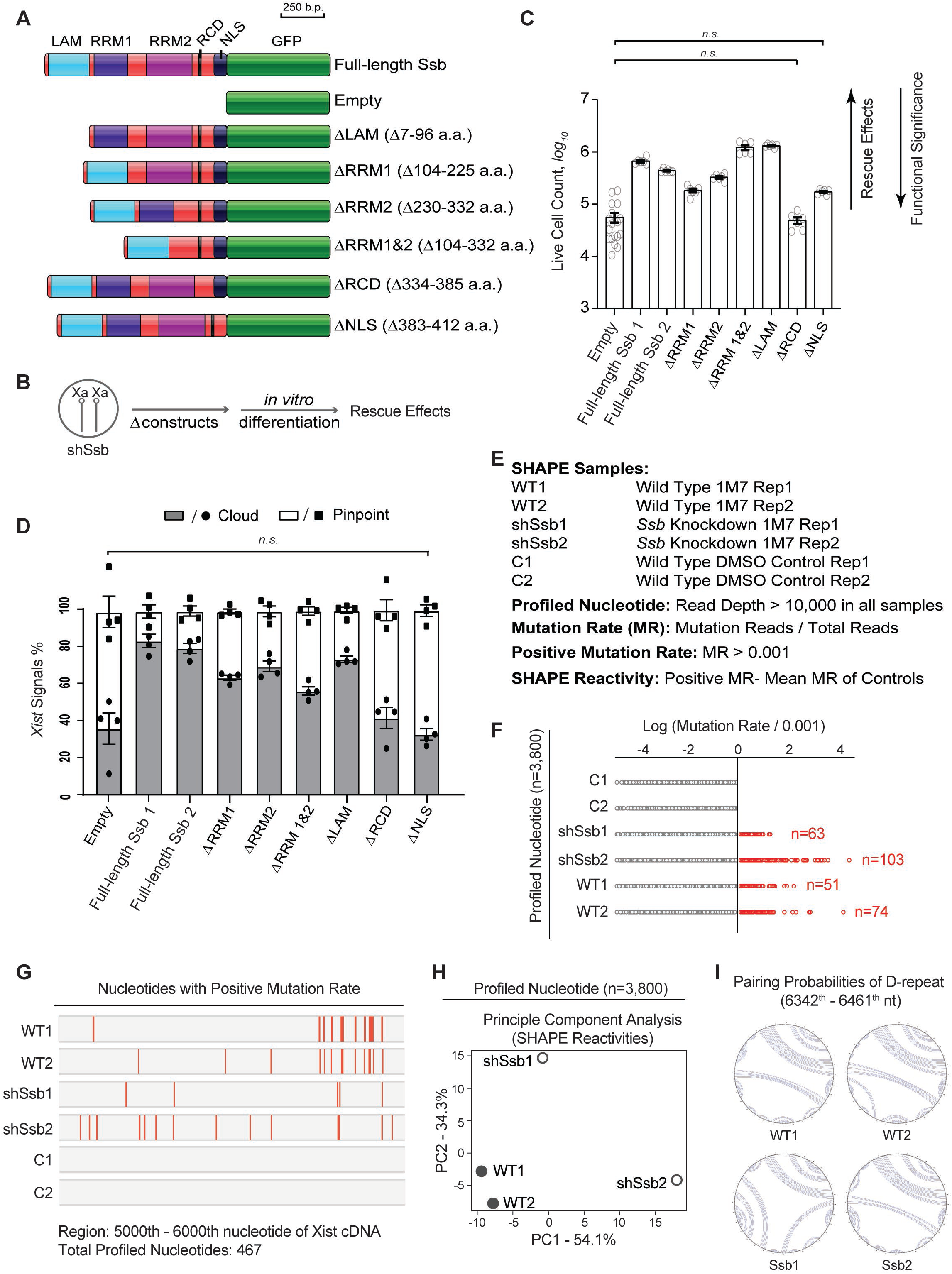
Dissecting the functional domains of La in XCI. **(A)** Diagram of the functional domains of La and the serial of plasmid constructs carrying in-frame deletions of selected functional domains (Δ constructs). **(B)** Experimental design of the rescue experiment. **(C)** The rescue effects of Δ constructs on cell survival of *Ssb* knockdown cells during *in vitro* differentiation. Cell counts of day 6 *in vitro* differentiation are shown. Data are shown as mean ± SEM. The statistical analysis used is the Student’s *t*-test. The data pairs with p > 0.05 (n.s.) are labeled. p < 0.0001 for all the rest of the pairwise comparison between Empty and any one of the serial Δ constructs. n = 18 from 6 independent experiments for Empty; n = 6 from 2 independent experiments for the rest of the constructs. **(D)** Quantification of *Xist* RNA FISH signals. Data are shown as mean ± SEM (n > 230 for all the constructs). Only the data pair with p > 0.05 (n.s.) is labeled. p < 0.0001 for all the rest of the pairwise comparison between Empty and any one of the serial Δ constructs. The statistical analysis used is *X*^*2*^ test. **(E)** Sample identities of SHAPE experiments and parameters used in data analysis. **(F)** Mutation rates of the 3,800 profiled nucleotides of each sample. **(G)** The distribution pattern of the profiled nucleotides with positive mutation rates along the region of 5000^th^ to 6000^th^ nucleotide of mouse *Xist* cDNA. Only profiled nucleotides with positive mutation rates are shown. The nucleotides are arranged in a sequential order based on nucleotide positions along *Xist*. Each nucleotide is represented by a red bar. **(H)** A PCA plot of the SHAPE reactivity profiles. **(I)** Pairing probabilities of D-repeat based on SHAPE reactivity profiles.

### *Xist* RNA is misfolded and less stable on *Ssb* knockdown background

We believe La is an *Xist*-binding protein important for folding of the RNA, possibly as an RNA chaperone. To assess the folding of *Xist* RNA, we performed SHAPE-MaP (selective 2’-hydroxyl acylation analyzed by primer extension and mutational profiling) (22). The RNA structure was probed by 1-methyl-7-nitroisatoic (1M7), which reacts with 2’-hydroxyl group and forms adducts along the RNA with a preference of unstructured regions. Chemical adducts along the treated RNA affect the fidelity of reverse transcription reaction and are detected as “mutation rates” by sequencing, which can be used to assess the RNA structure. Using male ES cell lines carrying the inducible *Xist* transgene, we performed in-cell SHAPE experiments in wild type cells (duplicate samples), *Ssb* knockdown cells (duplicate samples) and DMSO control (duplicate samples). Along the ∼18kb *Xist* RNA, we only analyzed the 3,800 nucleotides which were covered by more than 10,000 reads in each sample (“profiled nucleotides”, Fig. 6E and Table S5). We selected 0.1% mutation rate as the threshold. A mutation rate greater than 0.1% is considered “positive”. 0.1% is close to the error rate of the current Illumina sequencing technology and occurs to be a natural boundary separating the two control samples and the four 1M7-treated samples in our experiments.

Among the profiled nucleotides, all the 291 incidents of positive mutation rate were detected in 1M7-treated samples (Fig. 6F). The distribution patterns of the nucleotides with positive mutation rates are consistent between the two wild type samples. These data validate the SHAPE experimental system. Interestingly, the distribution patterns of the wild type samples are clearly distinguishable from mutation samples; and the two mutant sample also different from each other (Fig. 6 G-I). This observation can be best illustrated by the 467 profiled nucleotides along the 5000^th^-6000^th^ nucleotide region of *Xist* (Fig. 6G). Principle component analysis of the global SHAPE reactivity profiles and the structure predictions based on SHAPE reactivities are also in consistent with this notion (Fig. 6 H and I). These data show that *Xist* RNA is misfolded in mutant cells. Very likely, the misfolded RNA forms a random pool of transcripts which lacks a consensus structure and shows poor consistency from sample to sample.

We further investigated whether the misfolded *Xist* transcripts are subjected to degradation. To assess the stability of *Xist* RNA on *Ssb* knockdown background, we studied the disappearance of the *Xist* cloud signals after Dox removal (the “sunset” process). Cells were cultured in differentiating conditions and treated with Dox overnight before Dox removal. The rate of the sunset process reflects the stability of the *Xist* RNA. We observed a significantly faster sunset rate on *Ssb* knockdown background (Fig. 7 A and B), especially within the first two hours after Dox removal. To further assess the stability of *Xist* RNA, we performed quantitative RT-PCR to quantify the *Xist* RNA during sunset. The cell lines used were male ES cell lines carrying the inducible *Xist* transgene (Fig. 7 C and D). *Xist* transcription was blocked by Dox removal and actinomycin D treatment. The data was best fitted using a first-order exponential decay model (Fig. 7C). Half-life of *Xist* RNA was then calculated (Fig. 7D). Consistent with our live-cell imaging data, quantitative RT-PCR results show that the half-life of *Xist* is significantly shorter on *Ssb* knockdown background.

**Figure 7.**
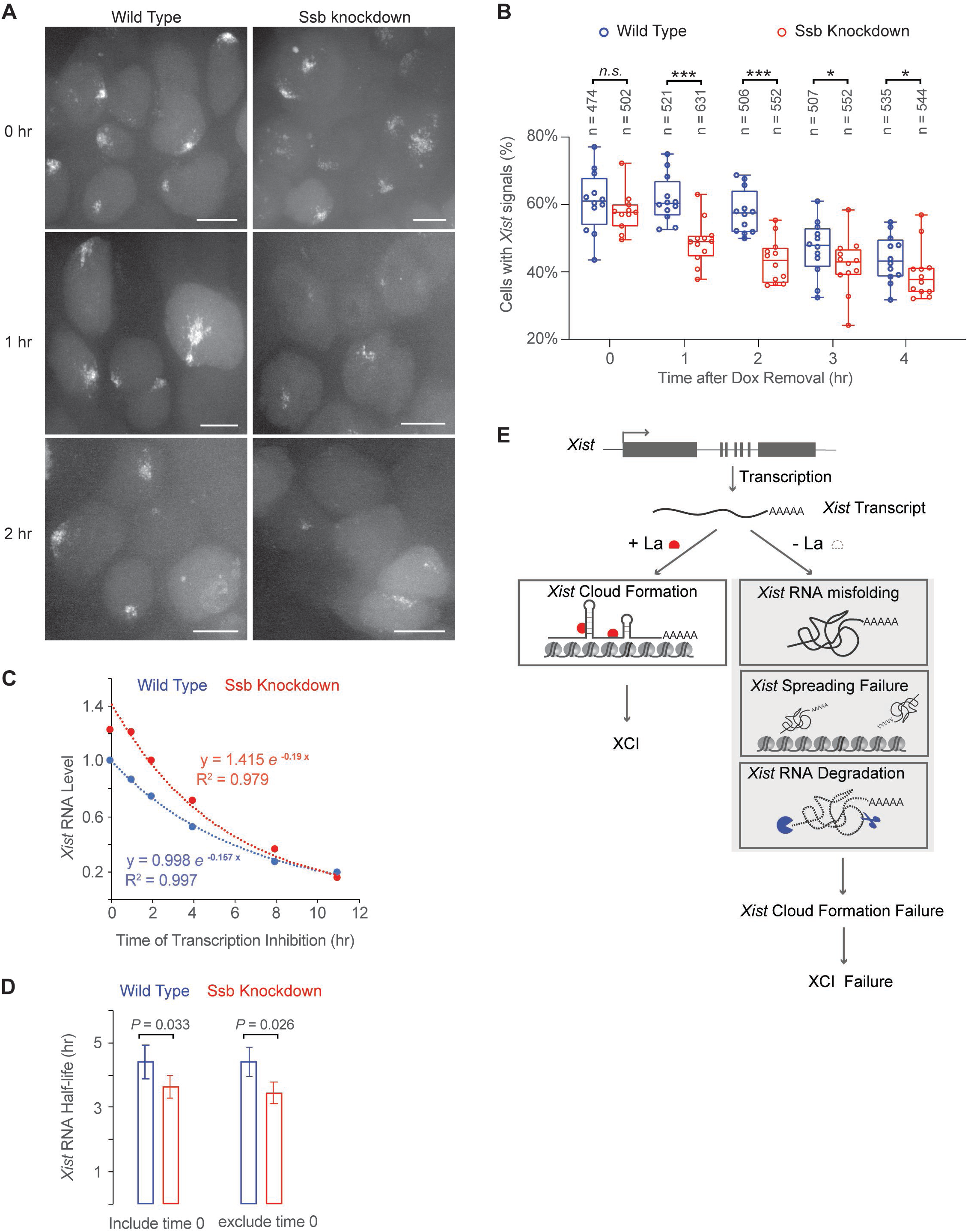
Stability of *Xist* on *Ssb* knockdown background by live-cell imaging after Dox removal, and a proposed model of how La is involved in XCI as an *Xist*-binding RNA chaperone. **(A and B)** The disappearance of *Xist* cloud signals in live cells after Dox removal. Cells were treated with Dox overnight in differentiating conditions. *Xist* cloud signals were assessed hourly after Dox removal. Representative images are shown. Scale bars, 8 μm. At each time point, images were collected from ∼12 randomly selected fields. The percentage of cells with *Xist* clouds in each field was quantified and the results are shown in box plots. The total number of cells analyzed at each time point is shown on the top. The statistical analysis used is the Student’s *t*-test. *p = 0.05; ***p < 0.01. **(C and D)** Measurement of *Xist* stability. Male ES cell lines carrying the inducible *Xist* transgene, with or without *Ssb* shRNA, were treated with Dox for 24 hours. Dox was then removed, and Actinomycin D (5 μg/ml) was added (defined as 0 time point). Quantitative RT-PCR was performed to measure *Xist* RNA levels in samples collected at indicated time points. Normalization was performed using *Actb*. Data was fitted using an exponential decay model with corresponding R-squared shown. The half-life of *Xist* was then calculated. Error bars indicate SD (n=3). **(E)** The proposed model.

Interestingly, it seems the decay reaction kinetics of *Xist* in *Ssb* knockdown cells can be separated into two phases (Fig. 7C). The first hour is the first phase of the reaction, which is a short phase and only contains two data points. The rest of the data points form the second phase of the decay reaction and can be better fitted in one reaction curve separately from the 0 time point. The half-life of *Xist* RNA estimated from the data of the second phase is shorter and with more statistical significance (Fig. 7D). We provide our interpretation on this in the discussion section.

Taken together, SHAPE results show that the *Xist* RNA transcripts are misfolded on *Ssb* knockdown background, and the shorter half-life of *Xist* RNA suggests that the misfolded *Xist* RNA transcripts are subjected to faster degradation.

## DISCUSSION

Here we show that FLAG-out is a useful method for profiling the interactome of lncRNAs. The advantage of FLAG-out is that once the target RNA is FLAG-tagged, the high specificity and high sensitivity of anti-FLAG antibody can be applied for the subsequent protein isolation. It should be noticed that selection of the PBS insertion site along the target RNA affects the outcome of FLAG-out. In this study, the PBSb sites are fused at the 5’ end of *Xist*, a position close to the most critical functional domain of *Xist*. In our previous study, we have shown that the 5’ PBSb fusion does not affect the *Xist* functionality (10).

La is known not only as an RNA chaperone, but also a helicase able to utilize ATP to unwind RNA-RNA and RNA-DNA duplex (14, 21, 23). Base on the results of this study, we propose that La is an *Xist*-binding protein involved in *Xist* RNA folding and spreading (Fig. 7E). In the mutant cells, the misfolded *Xist* RNA transcripts are unstable and unable to spread and form the *Xist* clouds. These defects eventually lead to failure of *Xist* cloud formation and collapse of XCI.

La is an autoimmune antigen found in the serum of patients with autoimmune diseases. More than 80% of the patients with autoimmune disorders are female (13). It is a possibility that XCI is related to the striking sex ratio distortion of autoimmune disorders. Multiple hypotheses have been developed. As numerous X-linked genes are related to immune functions, dysregulated expression of these genes may cause malfunction of the immune system. Therefore, skewed XCI pattern (loss of mosaicism), imbalanced X-linked gene dosage (X chromosome reactivation in early T cell lineage and haploinsufficiency in X chromosome monosomy) have all been hypothesized as possible causes of the sex ratio distortion in autoimmune disorders (13, 24). In *Ssb* knockdown cells, dysregulated gene expression occurs on many X-linked genes including the genes functionally related to immune functions (Fig. 2 F and G). These results suggest that *Ssb* may be a potential genetic factor of autoimmune diseases. However, identifying the lupus autoantigen La, as a protein involved in XCI does not help to further explain the sex ratio distortion. Rather, we believe it is a consequence of autoimmune disorders. Nuclear RNPs are immunogenic (25). The inactive X chromosome coated with *Xist* RNA and its associated proteins is a large piece of nuclear RNP in female cells, which can trigger autoimmune responses when exposed to the immune system under abnormal situations.

Understanding the structure of lncRNAs is critical for understanding lncRNAs’ functionality. Initial efforts to elucidate the *Xist* RNA structure have been reported, but significant inconsistency exists among the results of the initial studies (26-29). Nonetheless, several interesting and important issues are revealed. First, the in-cell structure of the RNA is significantly different from the *ex-vivo* structure of the RNA purified with mild conditions. Second, subregions of *Xist* may form dynamic structures in living cells. SHAPE analysis suggests that the dynamic structures function as loading pads for protein recruitment (27). In PARIS, a method in which the RNA structures are fixed in cells and the interacting regions of RNAs (duplex sequences) are directly sequenced, significant amount of conflicting duplex sequences is detected, which shows the *Xist* structure is highly dynamic in living cells (28). In our study, identifying La as a protein involved in *Xist* folding provides an important piece to the puzzle. Interestingly, La possesses three RRMs, which may function in a synergistic manner in regulating *Xist* structure in living cells. The mechanistic details await future research. Here, we discuss three possible ways to imagine the mechanistic roles of La in XCI. The three possibilities are not mutually exclusive to one another. Firstly, it is possible that La may function as an *Xist* chaperone. Indeed, the RCD domain of the human La protein has been characterized as an RNA chaperone domain (21). La proteins have also been characterized as RNA chaperones in different biological events (20). Interestingly, it has been hypothesized that RNA chaperones may function as “capacitors” to store and release the effect of genetic variation (20). Thus, the capacitor function of La may help to explain the structural conservation of *Xist*, a lncRNA with poor sequence homology among mammalian species (1). Secondly, as La is able to utilize ATP to unwind RNA-RNA and RNA-DNA duplex, it may play a more active role in controlling the structural dynamics of *Xist* in living cells (14, 23). These structural dynamics may be important for recruiting proteins onto the RNA and spreading of the RNA along its host chromosome territory. It should be noticed that the serial deletion approach used in this study cannot distinguish the RNA chaperon activity and the helicase activity of La, as the RCD domain (349-356 a.a.) and the ATP binding motif (348-355 a.a.) are largely overlapped (21, 23). Thirdly, La may be more directly involved in recruiting proteins onto *Xist*. Misfolding of *Xist* observed in mutant cells may be a consequence of protein loading failures.

As a protein involved in tRNA maturation, a significant fraction of La is located in the nucleolus (30, 31). The nucleolus may function as a general factory for assembling nuclear RNP particles, not just for rRNAs. *Xist* and its associated proteins forms a unique RNP complex in female cells. The fact that La is an *Xist*-binding RNA chaperone may be one reason behind the frequent association of the Xi with the nucleolus (32, 33).

It is an interesting observation that the decay reaction of *Xist* in *Ssb* knockdown cells seems to consist of two phases. The first hour of the reaction is a transient phase in which the decay reaction shows a slow kinetics. The rest of the data points form the second phase of the reaction which shows a significantly faster kinetics. To interpret the data, we assume that the decay reaction of *Xist* is catalyzed by both endonucleases and exonucleases. Endonuclease creates nicks along the RNA, which further serve as the starting points for exonucleases. Without the RNA chaperone activity of La, misfolded *Xist* RNA may be depleted for protein-binding and the RNA may be randomly folded in order to “protect” the hydrophobic bases from the hydrophilic environment. Therefore, compared to the protein-binding biologically active form, the misfolded structures may be more compact, which prevents the initial endonuclease reaction. Once the endonuclease reaction generates enough nicks to trigger the action of exonucleases, the degradation reaction quickly speeds up. However, we do not have enough data to study the transient first phase in detail, therefore, this interpretation may be speculative at this point. The details of the decay reaction and, more importantly, the structural details of *Xist* await future studies.

## MATERIALS AND METHODS

### Cell lines and culture

Mouse ES cells were cultured in medium containing 1000 unit/ml LIF. Feeder cells used were Drug Resistant 4 Mouse Embryonic Fibroblasts (DR4-MEF) (Applied StemCell; ASF-1002). Feeder-free ES cells were cultured on culture dishes pre-treated with 0.2 % gelatin (Sigma-Aldrich; G2500).

For *in vitro* differentiation, cells were cultured in differentiation medium containing 50 μg/ml L-ascorbic acid (Sigma). For the first 4 days of *in vitro* differentiation, embryoid bodies (EBs) were cultured in suspension. On day 4, EBs were transferred to a gelatinized T75 tissue culture flask. On day 5, EBs failed to attach to the tissue culture flask were washed away. On day 6, surviving cells were harvested for subsequent experiments.

Detailed information is provided in Supplementary Materials and Methods.

## ACKNOWLEDGEMENTS

L.C. was supported by the National Key R&D Program of China (Grant No. 2018YFA0107002 and 2018YFC1313003), the National Natural Science Foundation of China (Grant No. 31622038, 31671497 and 31871485), the Natural Science Foundation of Tianjin (Grant No. 18JCJQJC48400), the 111 Project Grant (B08011), and the Fundamental Research Funds for the Central Universities. L-F.Z. was supported by Singapore Ministry of Education Academic Research Fund (MOE2015-T2-1-093) and by the Singapore National Research Foundation under its Cooperative Basic Research Grant administered by the Singapore Ministry of Health’s National Medical Research Council (NMRC/CBRG/0092/2015).

