## Supplementary Materials for "The lupus autoantigen La is an *Xist*-binding protein involved in *Xist* folding and cloud formation"

**Author Contributions:** N.H., N.D., R.H., R.L., Y.L., X.D., X.W., and P.N. performed experiments, N.H., X.R., L-F.Z. and L.C. analyzed the data, L-F.Z. and L.C. designed the experiments and wrote the paper.

**This PDF file includes:**

Supplementary text
Supplementary Materials and Methods
Figures S1 to S8
Table S1
SI References

**Other supplementary materials for this manuscript include the following:**

Tables S2 to S5
Movies S1 to S7

**Supplementary Information text**

**Comparison of the candidate proteins of this study with previous studies:**

All 81 proteins identified by Chu et al (1) (annotated as C in Table S1) and all 10 proteins identified by McHugh et al (2) (annotated as N in Table S1) were compared with the candidate proteins identified in this study. The study of Minajigi et al (3) (annotated as S in Table S1) identified more than 700 proteins. The top 93 proteins ranked by "log2_avgFX/avgMX" were selected and compared with the candidate proteins identified in this study.

**Definition of functional groups:**

The candidate proteins are classified into each functional group based on the protein's function description in Uniport and NCBI gene.

Nuclear actin and related proteins:

1) The protein is myosin, unconventional myosin or myosin-binding.
2) The protein is actin-like, actin-related, actin-binding, actin filament binding, actin-crosslinking or regulates actin dynamics and organization.

Chromatin and related proteins:

1) The protein is involved in regulating chromatin structure, recruiting chromatin-remodeling enzymes, mediating nuclear import of histones and nucleosome assembly.
2) The protein possesses, recruits or regulates histone modification enzyme activities.
3) The protein binds to histones or modified histones.
4) The protein is a histone or histone variant.

DNA and RNA binding:

1) The protein is RNA-binding or DNA-binding.

Nuclear RNPs, ribosomal and nucleolar proteins:

1) The protein is a nuclear RNP.
2) The protein is a ribosomal protein.
3) The protein is a nucleolar protein.
4) The protein is involved in processing rRNA.

Membrane Proteins:

1) The protein is associated with plasma membrane, mitochondria membrane, trans-Golgi network and vesicle trafficking

**Supplementary Materials and Methods**

**FLAG-out**
Three 15 cm dishes of ES cells were cultured in differentiation medium supplemented with doxycycline for 48 hours. Cells were washed with ice cold PBS once and crosslinked with 0.4 J/cm$^2$ of $UV_{254\ nm}$. Cells were then harvested by trypsin treatment and lysed in 3 ml of lysis buffer (50 mM Tris HCl, pH 7.4; 150 mM NaCl; 1% TRITON X-100; 5% glycerol; supplemented with protease inhibitors and RNase Inhibitor). Cell lysates were first digested with TURBO DNase (Ambion). After centrifugation, supernatants were collected for co-IP experiment using anti-Flag M2 beads (Sigma). Elution of IPed proteins from anti-Flag M2 beads was achieved by 5 rounds of RNA/DNA digestion using 250 U of Benzonase (Millipore, #71206-3) for 2 h at 37°C. The eluted proteins were concentrated in a speed-vacuum to a final volume about 60 μl. One third of the elusions were separated on a 10% SDS-PAGE gel and stained by colloidal blue. The rest of the elusions were run on another 10% SDS-PAGE gel shortly, and the unresolved gel slices were subjected to mass spectrometry analysis. For proteins identified in both i-FLAG-*Xist* and i-Empty samples, ranking numbers were assigned according to the ranks of their protein scores in each sample (Table S2D). The ranking gains were calculated as: the rank number in i-Empty - the rank number in i-FLAG-*Xist*, similar to the method described in the previous publication (1).

**Cell Survival Assay**
Doxycycline treatment of 1 μg/ml was used throughout the study. G418 (ThermoFisher) treatment was carried out at 250 μg/ml. If ES cells were cultured as undifferentiated ES cells, Alkaline phosphatase (AP) staining (Vector Laboratories) was used at the end of the assay for counting the cells. After AP staining, milk is added to the well to provide a white background for picture taking, then the whole tissue culture plate is put onto a scanner to scan the image. ImageJ (4) was used to analyze AP staining data.

**shRNA knockdown**
An shRNA system (OligoEngine, pSUPER RNAi System) was used. The following shRNA sequences were designed against *Mybbp1a* (5'-CCGGAGTGTATTTGGTCATAT-3'), *Tardbp* (5'-GAATATGAAACCCAAGTGAAA-3'), *Ssb*-1(5'-GAGAAGATTTACACTTCCTTT-3'), and *Ssb*-2 (5'-GAACAGATCAAATTGGATGAA-3'). 300 μg/ml Hygromycin B (ThermoFisher; 10687010) was used to select stably transfected cell lines. Single surviving ES cell colonies were picked and the knockdown efficiency was validated using RT-qPCR.

**Quantitative RT-PCR**
Total RNA was isolated by TRIzol (Life technologies). cDNA was synthesized using iScript reverse transcription kit (170-8840, Bio-Rad). The real-time PCR was carried out on the CFX Connect real-time PCR system (Bio-Rad) using the SsoAdvanced Universal SYBR Green Supermix (Bio-Rad). The following PCR primers were used: Actb (F: 5'- ACTGCCGCATCCTCTTCCTC-3', R: 5'-CCGCTCGTTGCCAATAGTGA-3'); Gapdh (F: 5'-CCAATGTGTCCGTCGTGGAT-3', R: 5'-TGCCTGCTTCACCACCTTCT-3'); Mybbp1a (F: 5'-GCCGCGAGTTCTTGGACTT-3', R: 5'-ATCTCCGAATCATTGGGCCTT-3'); Tardbp (F: 5'- AATCAGGGTGGGTTTGGTAACA-3', R: 5'-GCTGGGTTAATGCTAAAAGCAC-3'); Ssb (F: 5'-ACCAAGCAGACCACTCCCTG-3', R: 5'-GGTGGCGTCAGTTGGGAAAC-3'); Cybb (F: 5'-TGTGGTTGGGGCTGAATGTC-3', R: 5'-CTGAGAAAGGAGAGCAGATTTCG-3'); Xiap (F: 5'-CGAGCTGGGTTTCTTTATACCG-3', R: 5'-GCAATTTGGGGATATTCTCCTGT-3'); Il2rg (F: 5'-CTCAGGCAACCAACCTCAC-3', R: 5'-GCTGGACAACAAATGTCTGGTAG-3'); Mecp2 (F: 5'-CAGGGAGGAAAAGTCAGAAGACC-3', R: 5'-AATGGTGGGCTGAAGGTTGTA-3'); Hprt (F: 5'-GATTAGCGATGATGAACCAGGTT-3', R: 5'-CCTCCCATCTCCTTCATGACA-3'); Gpc4 (F: 5'-GGCAGCTGGCACTAGTTTG-3', R: 5'-AACGGTGCTTGGGAGAGAG-3'); Neomycin (F: 5'-GGCTATTCGGCTATGACTGGGC-3', R: 5'-GCAGTTCATTCAGGGCACCG-3'); tdTomato (F: 5'-CCGACATCCCCGATTACAAGAAGC-3', R: 5'-TTGTAGATCAGCGTGCCGTC-3'); Xist E1 (F: 5'-CGGCCTCTAGTTTGTCCATT-3', R: 5'-GATGGCATGATGGAATTGAG-3'); Xist E7 (F: 5'-TCCTACCATCATCTGCTTTGA-3', R: 5'-CATCCGTTCACAAAGTCCAG-3'); GFP (F: 5'-GATGAAGAGCACCAAAGGCGC-3', R: 5'-CCGTTGTTGATGGCGTGCAG-3').

**Padlock probe SNP capture**
The padlock probe library design and padlock SNP capture procedure were performed as previously described (5). Briefly, each reaction was carried out in 20 μl volume containing 1 unit Ampligase

(A3210K, Epicenter), 1 unit Phusion High-Fidelity DNA Polymerase (M0530, New England BioLabs), 1 × Phusion High-Fidelity DNA Polymerase buffer, 10 nM dNTP. cDNA was generated using SuperScript III first-strand synthesis system (18080-051, Life Technologies). Two hundred nanograms of single-stranded cDNA and 2 pmol padlock probe were used in each reaction. Nicotinamide adenine dinucleotide (Sigma; N0632) was provided in each reaction at a final concentration of 0.5 mM.
The multiplexed sequencing libraries were PCR amplified in a real-time PCR system (CFX Connect, Bio-Rad) using the following primers: CA-2-RA.Miseq (5′-AATGATACGGCGACCACCGAGATCTATCGGCTACACGCCTATCGGGAAGCTGAAG-3′); CA-2-FA.Indx7Sol (5′-CAAGCAGAAGACGGCATACGAGATGATCTGCGGTCTGCCATCCGACGGTAGTGT-3′); CA-2-FA.Indx45Sol (5′-CAAGCAGAAGACGGCATACGAGATCGTAGTCGGTCTGCCATCCGACGGTAGTGT-3′); CA-2-FA.Indx76Sol (5′-CAAGCAGAAGACGGCATACGAGATAATAGGCGGTCTGCCATCCGACGGTAGTGT-3′); CA-2-FA.Indx91Sol (5′-CAAGCAGAAGACGGCATACGAGATACATCGCGGTCTGCCATCCGACGGTAGTGT-3′); CA-2-FA.Indx92Sol (5′-CAAGCAGAAGACGGCATACGAGATTCAAGTCGGTCTGCCATCCGACGGTAGTGT-3′); CA-2-FA.Indx93Sol (5′-CAAGCAGAAGACGGCATACGAGATATTGGCCGGTCTGCCATCCGACGGTAGTGT-3′). The following sequencing primers were used: Read1.Miseq (5′-ATCGGCTACACGCCTATCGGGAAGCTGAAG-3′); IndexRead (5′-ACACTACCGTCGGATGGCAGACCG-3′). Sequencing was carried out on an Illumina NextSeq system.

**ATAC-seq**

Trypsinized cells were pelleted and washed with PBS. A total of 55,000 cells were lysed using cold lysis buffer (10 mM Tris-Cl, pH 7.4, 10 mM NaCl, 3 mM $MgCl_2$, 0.1% (v/v) Igepal CA-630) for 3 mins and centrifuged at 500 ×*g* at 4 °C for 10 mins. Pellets were resuspended with transposition reaction mix (2.5 µL TD, 2.5 µL TDE1, 22.5 µL nuclease-free water) (Nextera DNA library preparation kit, Cat No. FC-121-1030) and incubated in Thermomixer at 37° for 30 min with 1000 rpm. Transposed DNA samples were purified with Qiagen MinElute PCR purification kit (Cat No.: 28006) and eluted with 20 µL elution buffer from Qiagen MinElute PCR purification kit. The purified transposed DNA fragments were amplified by PCR. Each reaction was carried out in 50 µL volume containing 20 µL of transposed DNA, 2.5 µL of 25 µM Ad1_noMX primer (6), 2.5 µL of 25 µM Ad2.* indexing primer (6), 25 µL of NEBNext High-Fidelity 2× PCR Master Mix (Cat. No. M0541) with the following cycles: 1 cycle: 5 min at 72 °C, 30 s at 98 °C and 5 cycles: 10 s at 98 °C, 30 s at 63 °C, 1 min at 72 °C. Quantitative PCR was carried out on the PCR products to calculate the additional PCR cycles required before reaching saturation. Each qPCR reaction was carried out in 15 µL volume containing 5 µL of partially PCR-amplified DNA, 4.41 µL of nuclease-free $H_2O$, 0.25 µL of 25 µM Ad1_noMX primer, 0.25 µL of 25 µM Ad2.* indexing primer, 0.09 µL of 100× SYBR Green I (Cat. No. S-7563), 5 µL of NEBNext High-Fidelity 2× PCR Master Mix (Cat. No. M0541) with the following cycles: 1 cycle: 30 s at 98 °C and 20 cycles: 10 s at 98 °C, 30 s at 63 °C, 1 min at 72 °C. The additional number of PCR cycles (*N*) is the cycle that corresponds to one-third of the maximum fluorescent intensity. After determining the additional PCR cycles required, the remaining 45 µL of transposed DNA was further amplified with the following cycles: 1 cycle: 30 s at 98 °C and *N* cycles: 10 s at 98 °C, 30 s at 63 °C, 1 min at 72 °C. ATAC-seq libraries were two-stage purified using AMPure XP beads: 0.5X and 1.3X. Libraries are eluted in 20 µL nuclease-free water. The size and pattern of the ATAC-seq libraries were assessed by 6% TBE-PAGE. The concentration of each library sample was measured by KAPA Library Quantification kit (Cat. No.: kk4824). All libraries were pooled and a final quality check was performed using Bioanalyzer High-Sensitivity DNA Analysis kit. Sequencing was carried out on Illumina HiSeq 4000 system.
The followings were the primers used in this study:
Ad1_noMX (5’-AATGATACGGCGACCACCGAGATCTACACTCGTCGGCAGCGTCAGATGTG-3’);
Ad2.1_TAAGGCGA(5’-CAAGCAGAAGACGGCATACGAGATTCGCCTTAGTCTCGTGGGCTCGGAGATGT-3’);
Ad2.2_CGTACTAG (5’-CAAGCAGAAGACGGCATACGAGATCTAGTACGGTCTCGTGGGCTCGGAGATGT-3’);
Ad2.3_AGGCAGAA (5’-CAAGCAGAAGACGGCATACGAGATTTCTGCCTGTCTCGTGGGCTCGGAGATGT-3’)

Ad2.4_TCCTGAGC (5'-
CAAGCAGAAGACGGCATACGAGATGCTCAGGAGTCTCGTGGGCTCGGAGATGT-3')
Ad2.5_GGACTCCT (5'-
CAAGCAGAAGACGGCATACGAGATAGGAGTCCGTCTCGTGGGCTCGGAGATGT-3')
Ad2.6_TAGGCATG (5'-
CAAGCAGAAGACGGCATACGAGATCATGCCTAGTCTCGTGGGCTCGGAGATGT-3');
Ad2.7_CTCTCTAC (5'-
CAAGCAGAAGACGGCATACGAGATGTAGAGAGGTCTCGTGGGCTCGGAGATGT-3');
Ad2.8_CAGAGAGG (5'-
CAAGCAGAAGACGGCATACGAGATCCTCTCTGGTCTCGTGGGCTCGGAGATGT-3');
Ad2.9_GCTACGCT (5'-
CAAGCAGAAGACGGCATACGAGATAGCGTAGCGTCTCGTGGGCTCGGAGATGT-3')
Ad2.10_CGAGGCTG (5'-
CAAGCAGAAGACGGCATACGAGATCAGCCTCGGTCTCGTGGGCTCGGAGATGT-3').

**RNA immunoprecipitation (RIP)**
Three 15-cm dishes of ES cells were harvested, crosslinked with 1% formaldehyde for 10 min at room temperature, and quenched with 0.125 M glycine for 5 min. After centrifugation, cell pellets were resuspended in lysis buffer (50 mM Tris-HCl pH 7.4, 150 mM NaCl, 1% TRITON X-100, 5% Glycerol, 1 mM PMSF, 1 mM DTT, 1.52 mg/ml protease inhibitor cocktail, 400 U/ml RNase Inhibitor). Incubate the cells for 30-60 minutes on a rotor at 4°C. Treat the lysate with RQ1 DNase (Promega M6101) at 37°C for 10 min. Insoluble material was removed by centrifugation, and the supernatant was incubated with anti-FLAG M2 beads (Sigma M8823) overnight at 4°C. The beads were washed with IP 200 buffer (20 mM Tris HCl pH 7.4, 200 mM NaCl, 1 mM EDTA, 0.3% TRITON X-100, 5% Glycerol) for five times. RNA-protein complexes were eluted from FLAG-beads by adding Proteinase K digestion buffer (50 mM Tris HCl pH 7.4, 10 mM EDTA, 50 mM NaCl, 0.5% SDS, 400 µg/ml Proteinase K (Roche 3115844001)), and incubated at 55 °C for 1 h. Then RNA was extracted using Trizol (Invitrogen). Quantitative RT-PCR was performed to quantify the amount of *Xist* and *Actin* RNA. The following PCR primers were used: *Xist*-1 (F: 5′- GGACATTTAGGTCGTACAGGAA-3′, R: 5′-CCAGGCTAACTCAGACATGAAG-3′); *Xist*-2 (F: 5′- TGCATACGGTCCTATTGCTAAA-3′, R: 5′-CCATGACTCTGGAAGTCAGTATG-3′); *Xist*-3 (F: 5′- CCTGCAAGGGATACCGTTTAT-3′, R: 5′-ATGAAAGGCGAAGGAGTATGG-3′); *Xist*-4 (F: 5′- AGAGTTGTTCCAGACCAATGA-3′, R: 5′-CTCACCTAAGCCCAAAGAAAGA-3′); *Xist*-5 (F: 5′- CCATGTGGTTGTTGGGATTTG-3′, R: 5′-TTACTCAGAAGGCTGGAGAGA-3′); *Actb* (F: 5'- ACTGCCGCATCCTCTTCCTC-3', R: 5'-CCGCTCGTTGCCAATAGTGA-3').

**SHAPE-MaP**
SHAPE-MaP was performed as described elsewhere (7). Briefly, ES cells were washed once with phosphate buffered saline, followed by adding 900 µl fresh medium and 100 µl of neat DMSO or 100 mM 1M7 in DMSO. After immediate mixing, cells were incubated at 37 °C for 5 minutes. Then cells were harvested, and RNA was purified using RNeasy Mini Kit (Qiagen). The mutational profiling (MaP) reverse transcription was carried out with SuperScript II reverse transcriptase (Life Technologies) in 1× MaP buffer (50 mM Tris (pH 8.0), 75 mM KCl, 6 mM $MnCl_2$, 10 mM DTT and 0.5 mM dNTPs), using random primers. The cDNAs were amplified by PCR using *Xist*-specific primers and Q5 high-fidelity DNA polymerase (NEB). The amplified DNA fragments were gel purified and pooled together. Construction of high-throughput sequencing libraries and sequencing were carried out by Novogene. The following primers were used for PCR amplification of *Xist* cDNA: *Xist*_-3 (F: 5'-TAGGCCATTTTAGCTATGACTGT-3', R: 5'-TTTGAACTCCCAGACCTCTTC-3'); *Xist*_-1 (F: 5'-GAGACATGGTCTCATAAAGCC-3', R: 5'-TGTGTGGAACCGAGGAAATA-3'); *Xist*_3 (F: 5'-AGGACTACTTAACGGGCTTA-3', R: 5'-AGGGTAATCAATCACCTGCA-3'); *Xist*_4 (F: 5'-CCCAGCATCCCTTTCCATTTC-3', R: 5'-AATTGCCAATGTGCTATGAG-3'); *Xist*_6 (F: 5'-GTCTCCTTGTGTTGTCTAATTCG-3', R: 5'-TTCTGGACCTATTGGGAAGGG-3'); *Xist*_8 (F: 5'-ACAAAAAGCTTACAGGCCACA-3', R: 5'-AATAGACACAAAGCAAGGAAG-3'); *Xist*_10 (F: 5'-TACTGAGGGTGATGAGTCTGT-3', R: 5'-TCAGCAATGTCATATCAAACAC-3'); *Xist*_11 (F: 5'-TCCATTGACCACTTTTCTGAATCAC-3', R: 5'-AAGATACTTGTCTTAAACATTCTGC-3'); *Xist*_12 (F: 5'-AGCAGAAAGAGGGTTGTACG-3', R: 5'-TGATGGAATTGAGAAAGGGCAC-3'); *Xist*_14 (F: 5'-GGTTCCTACCACTATGCCCTG-3', R: 5'-AAAACCCCATCCTTTATGCAA-3'); *Xist*_15 (F: 5'-TCACATGCTTTCTTATTTCAGCC-3', R: 5'-AGTTAACACTGTGCACATTTAC-3'); *Xist*_16 (F: 5'-CCCATCTATACCCCCTCCAT-3', R: 5'-GCAAGGGTAGTATTAGGACCTTGAG-3'); *Xist*_18 (F: 5'-CGTCTGATAGTGTGCTTTGC-3', R: 5'-GGCTTGGGATAGGTCTGAAA-3'); *Xist*_19 (F: 5'-

TGTTGGTGTTTGCTTGACTTCC-3’, R: 5’-AAACTTTAAGGACTCCAAAGTAAC-3’); *Xist*_20 (F: 5’-AGCGGACTGGATAAAAGCAAC-3’, R: 5’-CATCACAGTCTAATTCCATCCTG-3’); *Xist*_21 (F: 5’-GCTTGGTGGATGGAAATATGG-3’, R: 5’-CGTTATACCGCACCAAGAAC-3’).

**RNA FISH, immunostaining and immuno-RNA FISH**

RNA FISH, immunostaining and immuno-RNA FISH were carried out as previously described (8). Immunostaining for H3K27me3 was performed using a mouse monoclonal antibody (Abcam; ab6002; 1:500) with a secondary antibody conjugated with Alexa-647 (ThermoFisher; A-21236; 1:1000). Immunostaining for H2AK119ub was performed using a rabbit monoclonal antibody (Cell Signaling Technology; D27C4; 1:2000) with a secondary antibody conjugated with Alexa-647 (Abcam; ab150075; 1:1000). Immunostaining was followed by RNA FISH. The *Xist* RNA was detected with Sx9 probe, a P1 DNA construct containing a 40 kb genomic fragment covering the *Xist* gene. Nucleotide analogs used in probe labeling were Cy3-dUTP (Amersham, Cat# PA53022).

**Plasmid constructs**

A mouse *Ssb* cDNA clone was purchased from OriGene (MG206549) and the sequence of the cDNA was confirmed by Sanger sequencing (data not shown). A series of in-frame deletions of various functional domains of *Ssb* were generated by PCR amplification using Herculase II Fusion Enzyme (Agilent Technologies) followed by Gibson Assembly (NEB). The sequences of the cloned cDNA fragments were confirmed by sequencing (data not shown).

For rescue experiments, the plasmid constructs were transfected into a female 3F1 ES cells with *Ssb* stably knockdown. G418 (ThermoFisher) was used at 250 μg/ml to select for stably transfected cells. Single surviving colonies were picked and examined using RT-qPCR. The shRNA construct, which worked efficiently against *Ssb*, was not designed against the 3’ UTR of the RNA. Therefore, the shRNA is against some of the rescue plasmid constructs. Nonetheless, transfecting the *Ssb* knockdown cells with the rescue plasmids should compensate the effect of *Ssb* knockdown and serve as a rescue assay to study the functional domains of La.

**Microscopy and live-cell imaging**

Wide-field fluorescent microscopy work was carried out on an Eclipse Ti microscope (Nikon) with a digital camera (Clara Series model C01, Andor).

Live-cell imaging was performed on a CorrSight spinning disk confocal system (FEI Company) equipped with an Orca R2 CCD camera (Hamamatsu). 1 day before imaging, 800K feeder-free ES cells were seeded on fibronectin-coated glass-bottom dishes (MatTec Corp). Prior to live-cell imaging, cells were washed with 1x PBS and replaced with imaging medium composing complete medium for differentiating ES cells with DMEM substituted with FluoroBrite DMEM (ThermoFisher). 1 μg/ml of doxycycline was supplemented to the imaging medium to induce *Xist* expression. For live-cell time-lapse video recording, cells were placed into the microscope cell culture chamber heated to 37 °C at least 1 hr before imaging. Imaging was carried out in a closed chamber maintained at 37 °C with 5% CO2 and 90% humidity. A 488-nm laser line (iChrome MLE-LFA) was set at 100% laser power. Images were acquired using a PlanApo 63x/ 1.4 N.A. oil-immersion objective (Zeiss) (heated to 37 °C) with standard filter sets. The exposure time was set at 200 ms. All live-cell time-lapse video recording, unless explicitly stated otherwise, was carried out in a 2-hr time span with a 2-min time interval. For each time point, a 10-μm thick Z-stack with a 1-μm interval was collected. Autofocus system (Focus Clamp) was used to minimize out-of-focus throughout the recordings. Time-lapse imaging was started 1 hr after the addition of doxycycline for differentiating cells. All acquired images were processed and analyzed using ImageJ (4). Drift correction was applied to all time-lapse recordings.

In the “sunset” experiments, the snap-shot images of *Xist* signals in live cells were captured with an 800-ms exposure time at 100% laser power in 10-μm Z-stacks at 1-μm intervals.

**Data analysis**

For padlock SNP capture, we only selected the SNPs which were allelotyped in all 6 samples. A nucleotide position with a read count less than 10 was considered as undetected and its read count was set to 0. Read counts of all the SNPs from one gene were combined to calculate the allelotype of the gene. Genes known to escape XCI were removed. The allelotype of each gene is calculated as Log ((129 count +10) / (Cast count +10)). To avoid division by 0, we added a pseudocount of 10 to each read count. The sequencing data is available in sequence read archive (SRA, accession number PRJNA545157).

For ATAC-seq, we mapped 10 million reads from each sample onto the mouse genome (mm9) using bowtie2 (9). The sequencing depth of each sample was normalized to the total number of reads with a single best alignment position along the mouse genome outside of chromosome X. The alignment position of each read indicates the site of a transposon insertion event. The number of insertion sites along a given 1 mb region was counted and named as the “Cut Count” of the region. The sequencing data is available in SRA (accession number PRJNA545157).

For SHAPE-MaP, the data analysis software “ShapeMapper_v1.2” was downloaded from Dr. Kevin Weeks lab website (https://weeks.chem.unc.edu/software.html) (10). The recommended sample data set “shapemapper_example_data” (https://weeks.chem.unc.edu/software.html) was also downloaded for software test runs. The source code written in Python 2 was revised for Python 3 updates. The source code was also modified to calculate SHAPE reactivity using one control ($MutationRate^{1M7}$ - $MutationRate^{DMSO}$) instead of two controls ( ($MutationRate^{1M7}$ - $MutationRate^{DMSO}$) / $MutationRate^{Denatured}$ ). The following parameters were customly adjusted for the data analysis of this study: primerLength = 30; minDepth = 5,000; trimBothEnds = on. The rest parameters were kept the same as the default setting of the software. The software was used for counting mutation and generating the parameters used for RNA structure analysis. Other data analysis, such as profiled nucleotides, mutation rate profiles and principle component analysis, was performed using Perl and Python scripts built in-house. The in-cell SHAPE sequencing data is available in SRA (accession number PRJNA589511).

**Supplemental Figures**

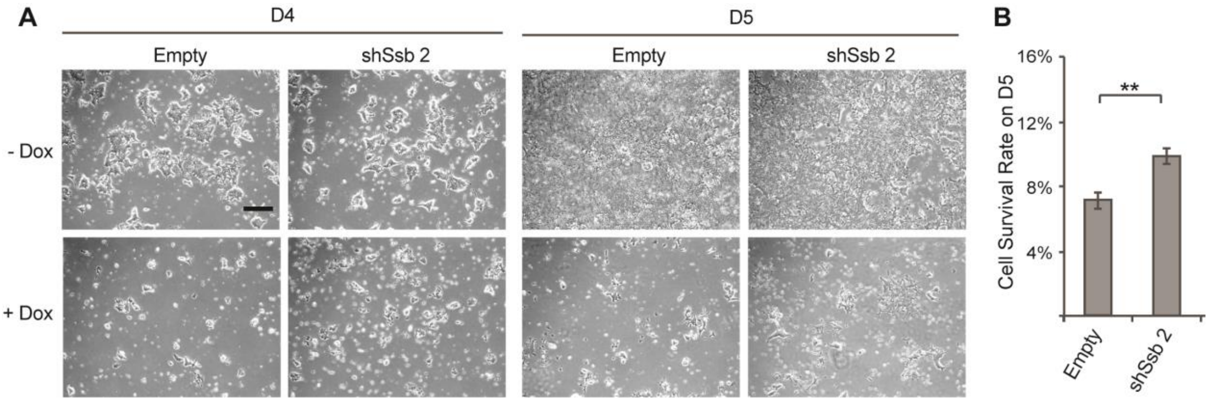

**Figure S1. The effects of *Ssb* knockdown on the induced XCI in differentiating male ES cells.** **(A)** Representative bright field microscope images of differentiating ES cell treated with Doxycycline for 4 and 5 days (D4 & D5). Clonal ES cell lines stably transfected with the shRNA constructs against *Ssb* and the empty shRNA vector are shown. Doxcycycline treatment was started upon *in vitro* differentiation. Scale bars, 300 μm. **(B)** Cell survival rate was calculated by cell counting on day 5. Data are shown as mean ± SEM. The statistical analysis used is the Student's *t*-test. **p < 0.01; n = 3.

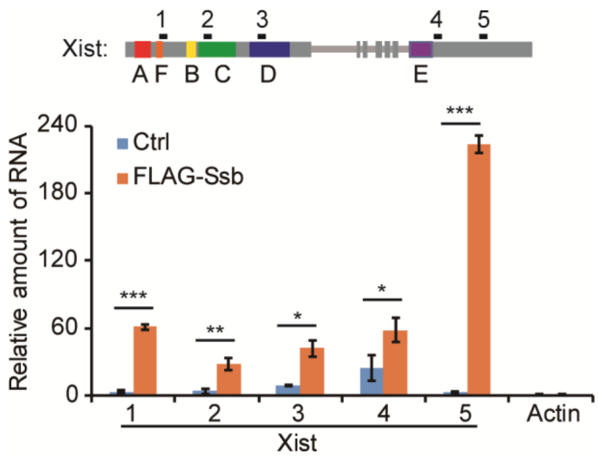

**Figure S2. RIP assay to validate the interaction between La and *Xist* RNA.** *Xist* inducible male ES cell lines with or without ectopic expression of FLAG-Ssb were treated with doxycycline treatment for 24 hours, and harvested for RIP experiments using anti-FLAG M2 beads. The top panel is a schematic illustration of the *Xist* locus. Repeat A-F are shown in coloured boxes. Amplicons of the PCR reactions are marked with short bars 1-5. The bottom panel shows the quantification results of RIP RNA. Normalization was performed using *Actb*. Error bars indicate SEM (n = 3). The statistical analysis used is the Student's *t*-test. *p < 0.05; **p < 0.01; ***p < 0.001.

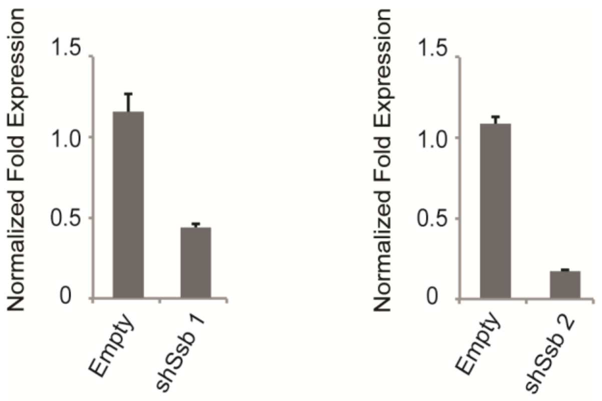

**Figure S3. Quantitative RT-PCR to assess the effect of shRNA knockdown of *Ssb* in the female 3F1 cell lines.** Data are shown in relative fold expression. Normalization was performed using *Gapdh*. Error bars indicate SEM (n = 3).

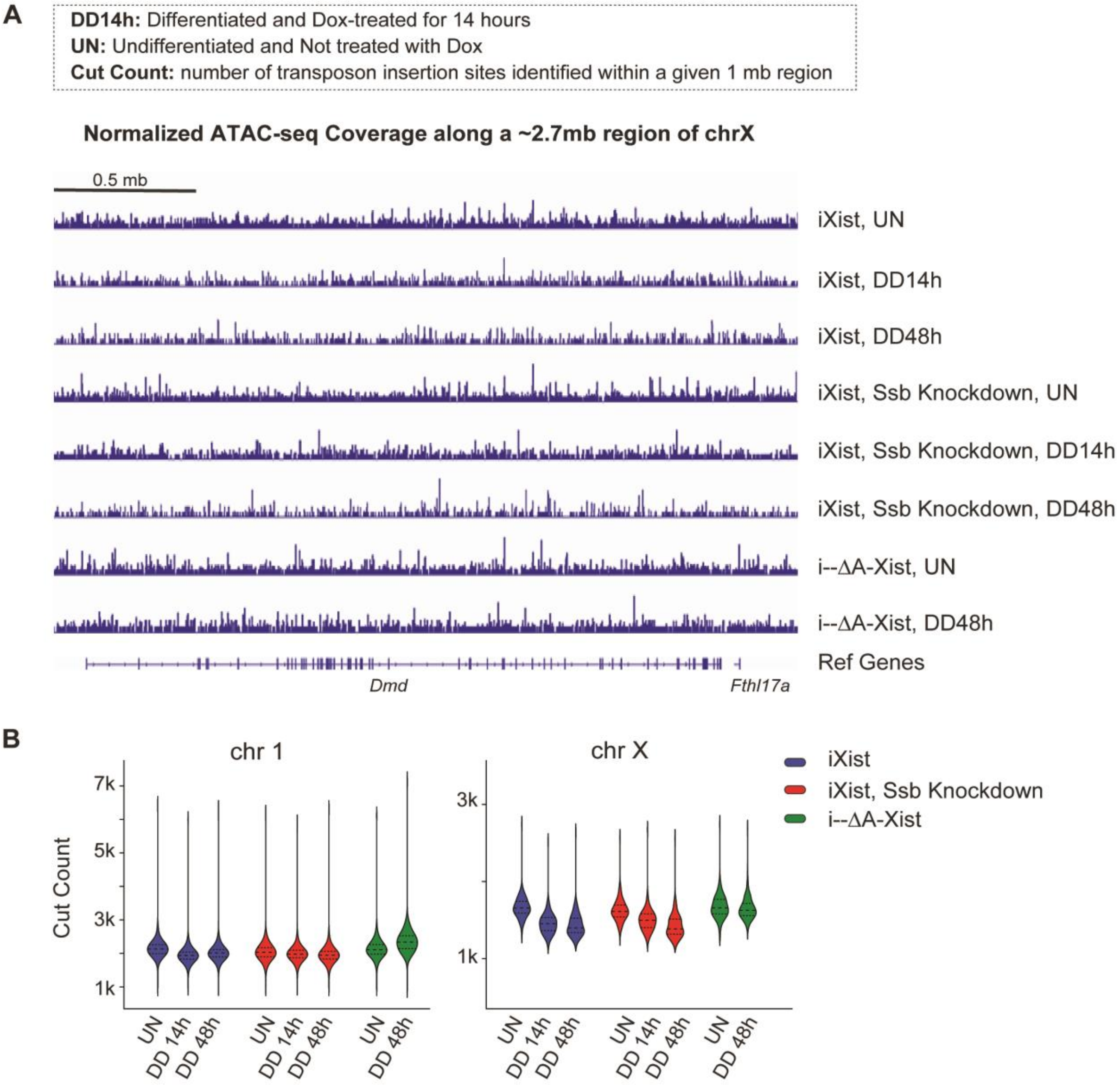

**Figure S4. ATAC-seq results. (A)** Normalized sequencing coverage along a ~2.7 Mb X-linked region. **(B)** The chromosomes are divided into 1-mb regions and the Cut Counts of each 1-Mb region are shown in violin plots.

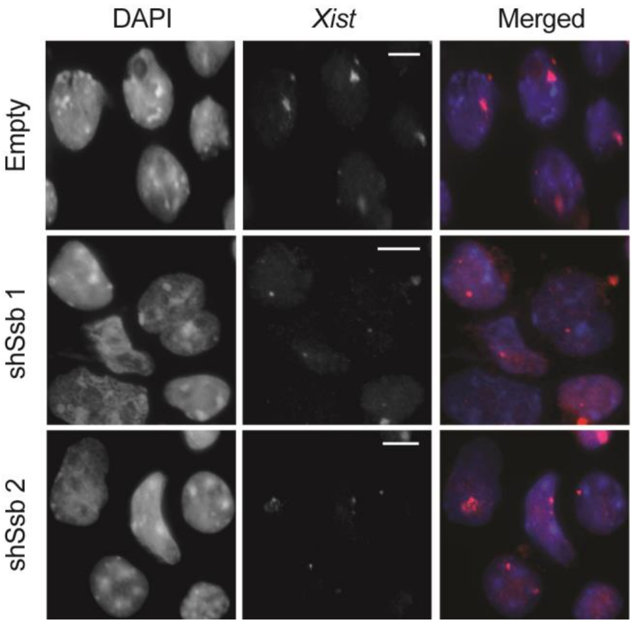

**Figure S5. *Xist* RNA FISH results.** The experiment was performed using day 6 *in vitro* differentiating female ES cells. DNA was counter stained with DAPI (blue). Scale bars, 8 μm.

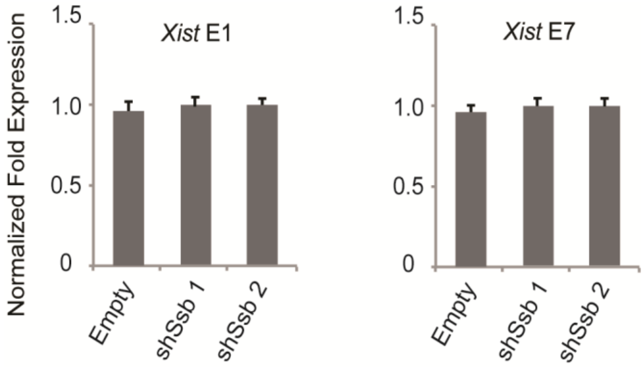

**Figure S6. Quantitative RT-PCR to assess the expressions of *Xist* RNA in the *Xist* inducible male ES cell line upon *Ssb* knock-down.** Cells were cultured as ES cells and Doxycycline treatment was carried out for 14 hours. Primers used were either targeting *Xist* Exon 1 (E1) or Exon 7 (E7). Data are shown in relative fold expression. Normalization was performed using *Gapdh*. Error bars indicate SEM (n = 6 for Empty; n = 3 for shSsb 1 and ShSsb 2).

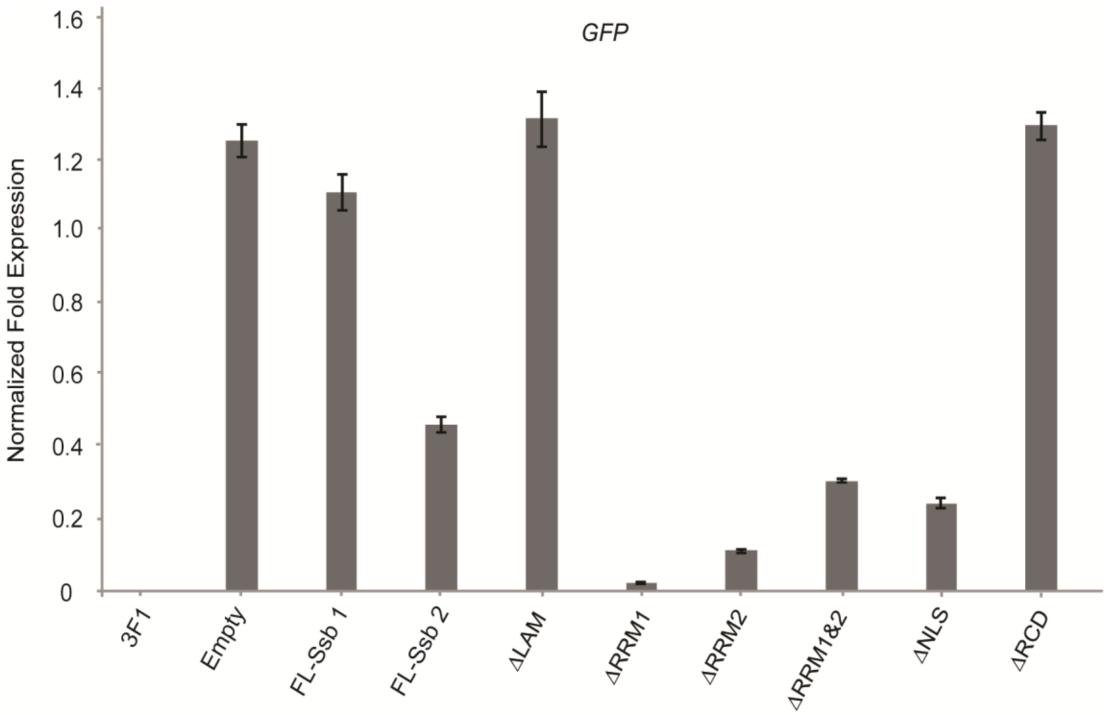

**Figure S7. Quantitative RT-PCR to assess the expression level of the GFP-fusion genes in the stable rescue lines.** Data are shown in relative fold expression. Normalization was performed using *Gapdh*. Error bars indicate SEM (n = 3).

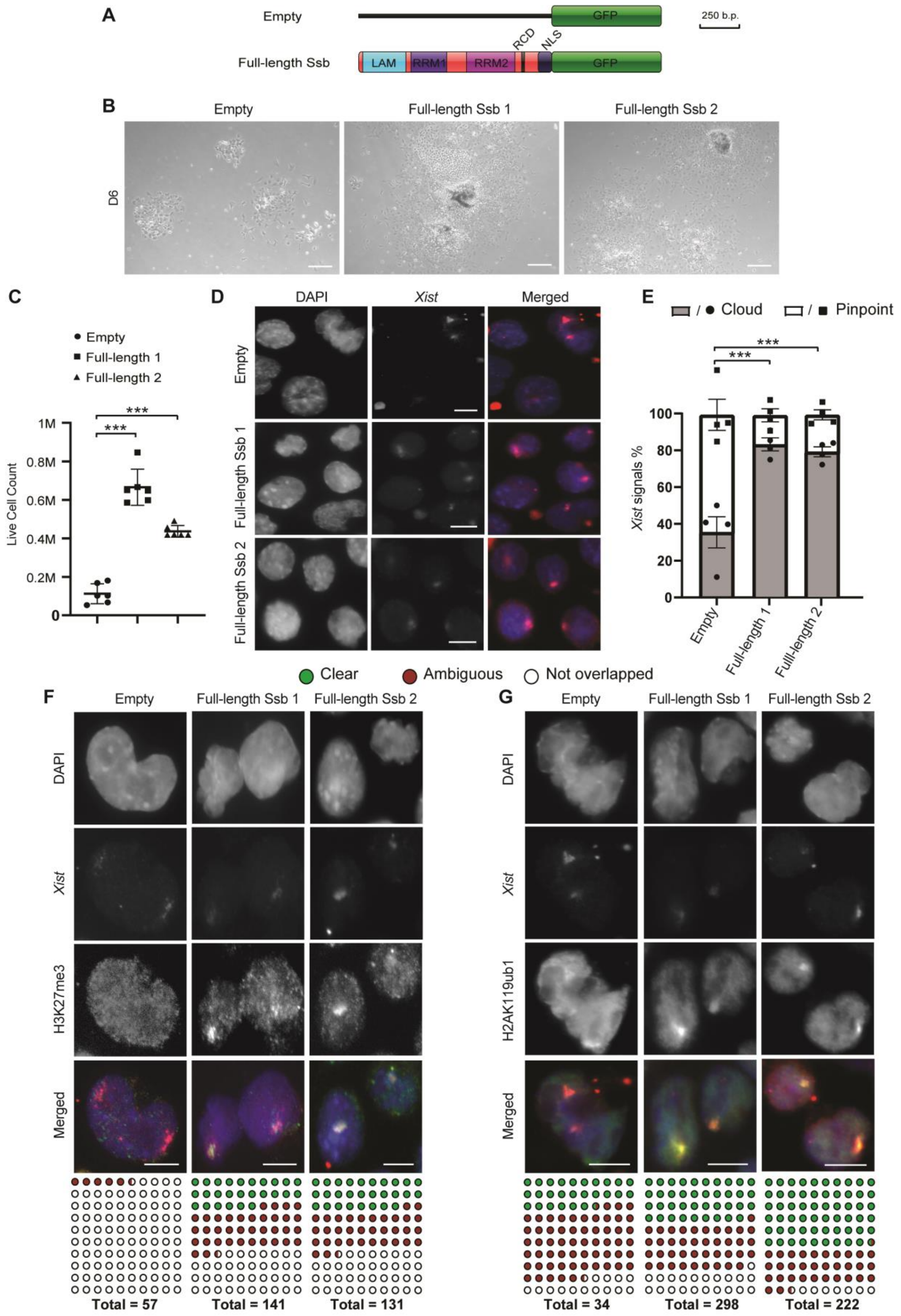

**Figure S8. The rescue effects of full-length *Ssb* constructs on *Ssb* knockdown female cells during *in vitro* differentiation. (A)** Diagram of the functional domains of La and the plasmid constructs. **(B)** Representative microscope images of day 6 differentiating ES cells. Scale bars, 300

μm. Cells are clonal ES cell lines stably transfected with the corresponding plasmid constructs. **(C)** The rescue effects of full-length *Ssb* constructs on cell survival of *Ssb* knockdown cells during *in vitro* differentiation. Cell counts of day 6 *in vitro* differentiation are shown. Data are shown as mean ± SEM (n = 6). The statistical analysis used is the Student's *t*-test. The data pairs with p > 0.05 (n.s.) are labeled. ***p < 0.001. **(D)** *Xist* RNA FISH. DNA was counterstained with DAPI. Scale bars, 8μm. **(E)** Quantification of *Xist* RNA FISH signals. Data are shown as mean ± SEM (n = 243 for Empty; n = 545 for Full-length 1; n = 839 for Full-length 2). ***p < 0.001 by $X^2$ test. **(F and G)** Immuno-RNA FISH detecting H2AK119ub, H3K27me3 and *Xist.* Cells were *in vitro* differentiated for 6 days. Immunostains were performed before the RNA FISH. DNA was counter stained with DAPI (blue). Scale bars, 8 μm. The total number of *Xist* clouds with clear, ambiguous or undetectable overlapping with the histone mark enrichment were tallied and tabulated below.

**Table S1** *Xist*-binding proteins identified by “FLAG-out”

| Nuclear Actin and Related Proteins | | |
|---|---|---|
| ID | Name | Also Identified in |
| Q8VDD5 | Myosin-9 | |
| Q8BFZ3 | Beta-actin-like protein 2 | |
| Q9QXZ0 | Microtubule-actin cross-linking factor 1 | |
| Q80X90 | Filamin-B | |
| Q9JI91 | Alpha-actinin-2 | |
| Q6URW6 | Myosin-14 | |
| Q9JMH9 | Unconventional myosin-XVIIIa | |
| Q9JKF1 | Ras GTPase-activating-like protein IQGAP1 | |
| Q91Z67 | SLIT-ROBO Rho GTPase-activating protein 2 | |
| Q5NBX1 | Protein cordon-bleu | |
| E9Q634 | Unconventional myosin-Ie | |
| P59242 | Cingulin | |
| P46735 | Unconventional myosin-Ib | |
| Q8BQ30 | Phostensin | |
| Q6R891 | Neurabin-2 | |
| Q8CI43 | Myosin light chain 6B | |
| Q9Z2N8 | Actin-like protein 6A | |
| Q9QXS6 | Drebrin | |
| Q9R0P5 | Destrin | |
| Q6P9R2 | Serine/threonine-protein kinase OSR1 | |
| Q61553 | Fascin | |
| Q9WTI7 | Unconventional myosin-Ic | |
| Q61879 | Myosin-10 | |
| Q8BTM8 | Filamin-A | |
| Q64331 | Unconventional myosin-VI | |
| Q9DBR7 | Protein phosphatase 1 regulatory subunit 12A | |
| Q9ERG0 | LIM domain and actin-binding protein 1 | |
| Q9QYC0 | Alpha-adducin | |
| Q9CVB6 | Actin-related protein 2/3 complex subunit 2 | |
| Q99JY9 | Actin-related protein 3 | |
| P14602 | Heat shock protein beta-1 | |
| Q2KN98 | Cytospin-A | |
| P58774 | Tropomyosin beta chain | |
| Q9QYB5 | Gamma-adducin | |
| Q3THE2 | Myosin regulatory light chain 12B | |
| O88990 | Alpha-actinin-3 | |
| O70318 | Band 4.1-like protein 2 | |
| Q9JJ28 | Protein flightless-1 homolog | |

| Chromatin and Related Proteins | | |
|---|---|---|
| ID | Name | Also Identified in |
| P62141 | Serine/threonine-protein phosphatase PP1-beta catalytic subunit | |
| Q7TPV4 | Myb-binding protein 1A | C, S |
| Q78ZA7 | Nucleosome assembly protein 1-like 4 | |
| O35129 | Prohibitin-2 | |
| O09106 | Histone deacetylase 1 | |

| | |
|---|---|
| Q99LL5 | Periodic tryptophan protein 1 homolog |
| P10853 | Histone H2B type 1-F/J/L |
| Q99J09 | Methylosome protein 50 |
| Q60972 | Histone-binding protein RBBP4 |
| Q62318 | Transcription intermediary factor 1-beta |
| P02301 | Histone H3.3C |
| Q9WTM5 | RuvB-like 2 |
| P70168 | Importin subunit beta-1 |

| **DNA and RNA Binding Proteins** | | |
|---|---|---|
| **ID** | **Name** | **Also Identified in** |
| P17225 | Polypyrimidine tract-binding protein 1 | C, N |
| Q60817 | Nascent polypeptide-associated complex subunit alpha | |
| Q8R4U7 | Leucine zipper protein 1 | |
| Q80U78 | Pumilio homolog 1 | |
| P11031 | Activated RNA polymerase II transcriptional coactivator p15 | |
| Q9CY58 | Plasminogen activator inhibitor 1 RNA-binding protein | |
| Q921F2 | TAR DNA-binding protein 43 | C, S |
| P53996 | Cellular nucleic acid-binding protein | |
| Q921M3 | Splicing factor 3B subunit 3 | |
| Q9DBD5 | Proline-, glutamic acid- and leucine-rich protein 1 | |
| P26369 | Splicing factor U2AF 65 kDa subunit | |
| Q8K3Y3 | Protein lin-28 homolog A | C |
| Q9CQF3 | Cleavage and polyadenylation specificity factor subunit 5 | |
| Q3U1J4 | DNA damage-binding protein 1 | |
| Q99KP6 | Pre-mRNA-processing factor 19 | |
| Q60865 | Caprin-1 | |
| Q02248 | Catenin beta-1 | |
| Q8BG81 | Polymerase delta-interacting protein 3 | C |
| P97855 | Ras GTPase-activating protein-binding protein 1 | |
| P32067 | Lupus La protein homolog | C |

| **Membrane Proteins** | | |
|---|---|---|
| **ID** | **Name** | **Also Identified in** |
| P39447 | Tight junction protein ZO-1 | |
| Q9Z0U1 | Tight junction protein ZO-2 | |
| P48962 | ADP/ATP translocase 1 | |
| Q68FD5 | Clathrin heavy chain 1 | |
| P23242 | Gap junction alpha-1 protein | |
| Q9WVE8 | Protein kinase C and casein kinase substrate in neurons protein 2 | |
| P17809 | Solute carrier family 2, facilitated glucose transporter member 1 | |
| Q7TMK6 | Protein Hook homolog 2 | |

| **Nuclear RNP; Ribosomal or Nucleolar Proteins** | | |
|---|---|---|
| **ID** | **Name** | **Also Identified in** |
| P47962 | 60S ribosomal protein L5 | |
| P35980 | 60S ribosomal protein L18 | |

| | | |
|---|---|---|
| P62911 | 60S ribosomal protein L32 | |
| P62267 | 40S ribosomal protein S23 | |
| O09167 | 60S ribosomal protein L21 | |
| P62862 | 40S ribosomal protein S30 | |
| P62717 | 60S ribosomal protein L18a | |
| P12970 | 60S ribosomal protein L7a | |
| P41105 | 60S ribosomal protein L28 | |
| P47911 | 60S ribosomal protein L6 | |
| P62855 | 40S ribosomal protein S26 | |
| P61514 | 60S ribosomal protein L37a | |
| P47915 | 60S ribosomal protein L29 | |
| P14148 | 60S ribosomal protein L7 | |
| P53026 | 60S ribosomal protein L10a | |
| Q9D8E6 | 60S ribosomal protein L4 | |
| P62242 | 40S ribosomal protein S8 | |
| P14206 | 40S ribosomal protein SA | |
| P47963 | 60S ribosomal protein L13 | |
| P63276 | 40S ribosomal protein S17 | |
| P47964 | 60S ribosomal protein L36 | |
| P62892 | 60S ribosomal protein L39 | |
| Q9D0E1 | Heterogeneous nuclear ribonucleoprotein M | C, N, S |
| Q9Z2X1 | Heterogeneous nuclear ribonucleoprotein F | |
| Q60668 | Heterogeneous nuclear ribonucleoprotein D0 | C |
| Q8VEK3 | Heterogeneous nuclear ribonucleoprotein U | C, N, S |
| Q61937 | Nucleophosmin | |
| Q9CPP0 | Nucleoplasmin-3 | |
| O54825 | Bystin | |
| Q62189 | U1 small nuclear ribonucleoprotein A | |

| **Other Proteins** | | |
|---|---|---|
| **ID** | **Name** | **Also Identified in** |
| O70251 | Elongation factor 1-beta | |
| O54931 | A-kinase anchor protein 2 | |
| Q9D8W5 | 26S proteasome non-ATPase regulatory subunit 12 | |
| O88844 | Isocitrate dehydrogenase [NADP] cytoplasmic | |
| O88735 | Ensconsin | |
| Q9R0Q9 | Mannose-P-dolichol utilization defect 1 protein | |
| Q61316 | Heat shock 70 kDa protein 4 | |
| Q9JKV1 | Proteasomal ubiquitin receptor ADRM1 | |
| Q8BFR5 | Elongation factor Tu, mitochondrial | |
| Q9CZS1 | Aldehyde dehydrogenase X, mitochondrial | |
| Q8BJM7 | S-adenosyl-L-methionine-dependent tRNA 4-demethylwyosine synthase | |
| P45376 | Aldose reductase | |
| P23116 | Eukaryotic translation initiation factor 3 subunit A | |
| P26516 | 26S proteasome non-ATPase regulatory subunit 7 | |
| P17751 | Triosephosphate isomerase | |
| O88685 | 26S protease regulatory subunit 6A | |
| P05064 | Fructose-bisphosphate aldolase A | |
| P68040 | Guanine nucleotide-binding protein subunit beta-2-like 1 | |

| | |
|---|---|
| Q9D4J1 | EF-hand domain-containing protein D1 |
| Q8QZY1 | Eukaryotic translation initiation factor 3 subunit L |
| Q8JZQ9 | Eukaryotic translation initiation factor 3 subunit B |
| P60229 | Eukaryotic translation initiation factor 3 subunit E |
| P35700 | Peroxiredoxin-1 |
| P17182 | Alpha-enolase |
| Q91VJ4 | Serine/threonine-protein kinase 38 |
| P24369 | Peptidyl-prolyl cis-trans isomerase B |
| Q3TXS7 | 26S proteasome non-ATPase regulatory subunit 1 |
| P99024 | Tubulin beta-5 chain |
| P01868 | Ig gamma-1 chain C region secreted form |

**Movies S1-3:** The emergence of induced *Xist* RNA signals in wild type differentiating ES cells.
**Movies S4-7:** The emergence of induced *Xist* RNA signals in *Ssb* mutant differentiating ES cells.
